# ISGylation Mechanism Uncovers Conformational Specificity for HECT-family E3 ligase

**DOI:** 10.1101/2025.09.26.678755

**Authors:** Pritiranjan Sahoo, Gloria Grace Parmar, Dipti Ranjan Lenka, Mohini Sherawat, Bhagya Shri Trivedi, Atul Kumar

## Abstract

Interferon-stimulated gene 15 (ISG15), composed of N-terminal and C-terminal ubiquitin-like domains (NTD/CTD), plays a critical role in antiviral immunity. Although the ubiquitination mechanism is well established, the mechanisms governing ISG15 transfer, particularly from E2 to E3 and subsequent lysine conjugation, remain unknown. Here, we reveal that UbcH8(E2)∼ISG15 exhibits striking specificity for HECT-family E3 ligases (particularly HERC5) but is inactive with RING or RBR E3. In contrast, UbcH8∼Ub preferentially engages RBR E3, indicating a switched E2–E3 specificity depending on the conjugated ubiquitin-like modifier. Structural and biochemical studies uncover how a unique closed conformation of UbcH8∼ISG15 enables trans-thiolation mediated by selective HECT-family E3 ligases. We further demonstrate that HERC5’s C-lobe specifically recognizes donor ISG15 for lysine conjugation, explaining its exclusive ISGylation activity and lack of ubiquitination function. These findings delineate the molecular basis of ISG15 conjugation and reveal how its pathway has evolved distinct mechanisms from ubiquitination, offering new avenues for therapeutic intervention in infection and immunity.

## Introduction

Type I interferons (IFNs-α/β) induce the expression of interferon-stimulated gene 15 (ISG15), a ubiquitin-like modifier best known for its antiviral functions^1, 2^. Beyond its role in host defense, emerging evidence implicates ISG15 in diverse physiological processes, including DNA replication, bacterial infection responses, and the regulation of female fertility through ovulation suppression^3–5^. During pathogen invasion, host cells secrete IFNs and proinflammatory cytokine s, triggering the upregulation of ISG15 and other effector proteins as part of the innate immune response^6^.

ISG15 conjugation (ISGylation) of host immune proteins can modulate their activity, as exemplified by its role in activating the viral sensor MDA5^7^. Beyond its immunomodulatory functions, ISG15 also directly targets viral proteins, inhibiting their activity and amplifying antiviral responses. For example, modification of the E3 ligase NEDD4 by ISG15 leads to its sequestration, bolstering host antiviral responses^8^. In turn, viruses have evolved countermeasures to evade ISG15-mediated restriction. For instance, SARS-CoV-1 and SARS-CoV-2 encode a papain-like protease (PLpro) that cleaves ISG15 from host and viral proteins, effectively reversing ISGylation^9, 10^. Similarly, Influenza B virus employs its NS1B protein to sequester ISG15, relocalizing it from the cytoplasm to nuclear compartments and disrupting ISGylation of nascent polyribosome-associated proteins^11–13^. NS1B also reduces the secretion of free ISG15, which functions as a cytokine-like mediator of inflammation^14, 15^. Notably, this interaction is species-specific: while human ISG15 binds NS1B, mouse ISG15 cannot^16–18^, potentially explaining Influenza B’s human tropism and implicating ISG15 in viral reassortment events that drive pandemic emergence.

ISG15 is a 17.8 kDa protein comprising two ubiquitin-like domains (NTD and CTD) that share ∼30% sequence similarity with ubiquitin (Ub) while maintaining nearly identical structural folds (RMSD <1.8 Å)^19, 20^. Like other ubiquitin-like modifiers (Ubls), ISG15 is conjugated to target proteins via a three-enzyme cascade: the E1 activating enzyme (UBA7/UBE1L, ISG15-specific)^21^, the E2 conjugating enzyme UBE2L6/UbcH8 (shared with ubiquitination)^22–26^, and E3 ligases. HERC5 (HECT family), HHARI (RBR family), and TRIM25 (RING family) have been proposed as major ISG15 E3 ligases ^24, 27–29^. However, to date, the catalytic mechanism of E3-mediated ISGylation remains unknown. Also, while the activity of Parkin (RBR family) and NEDD4 (HECT family) E3 ligases is affected by ISGylation^8, 30^, it is not known whether these two have any role in protein ISGylation as E3 ligases. Mechanistically, recent cryo-EM studies have illuminated the UBE1L-UbcH8 interaction^31, 32^. However, the structural basis for ISG15 transfer from UbcH8 to E3 ligases and subsequent conjugation to lysine residues remains unknown, representing a critical gap in understanding ISGylation specificity.

Based on the catalytic mechanism, E3 ligases are classified into three major classes: RING, HECT, and RBR family E3 ligases. Ubiquitin (Ub) adopts distinct conformational states during E2∼Ub recognition and transfer to E3 ligases, dictating specificity in ubiquitination (Fig. 1a, Extended Data Fig. 1)^33–35^. The latter underlies a conserved catalytic mechanism among different members of the same family of E3 ligases, despite their varying substrates. Ub maintains a closed conformation (I44 patch buried against the E2 crossover helix) with RING-family E3s, enabling direct Ub transfer from the E2 catalytic cysteine to lysines (Fig. 1a, Extended Data Fig. 1)^36^. In contrast, RBR-family E3s stabilize an open Ub conformation (I44 patch engaged by the E3), facilitating trans-thiolation to form the E3∼Ub intermediate (Fig. 1a, Extended Data Fig. 1)^37^. While HECT-family E3s were initially thought to exclusively promote an open Ub state (I44 solvent-exposed)^38^ (Fig. 1a, Extended Data Fig. 1), recent cryo-EM structure reveal HECT induce a closed E2∼Ub conformation essential for trans-thiolation^39^. Following the trans-thiolated state of E3∼Ub, interactions between the donor ubiquitin (linked to the catalytic cysteine of HECT and RBR E3s) and the E3 ligase (in both HECT and RBR families) regulate the subsequent transfer of ubiquitin to lysines (Fig. 1a). Notably, while these Ub conformational dynamics are well-characterized, the structural mechanisms governing ISG15 transfer, particularly during E2-E3 trans-thiolation and final lysine conjugation, remain undefined. Given that several E2/E3 enzymes participate in both ubiquitination and ISGylation, elucidating ISG15-specific conformational changes is critical for understanding pathway selectivity.

**Figure 1.**
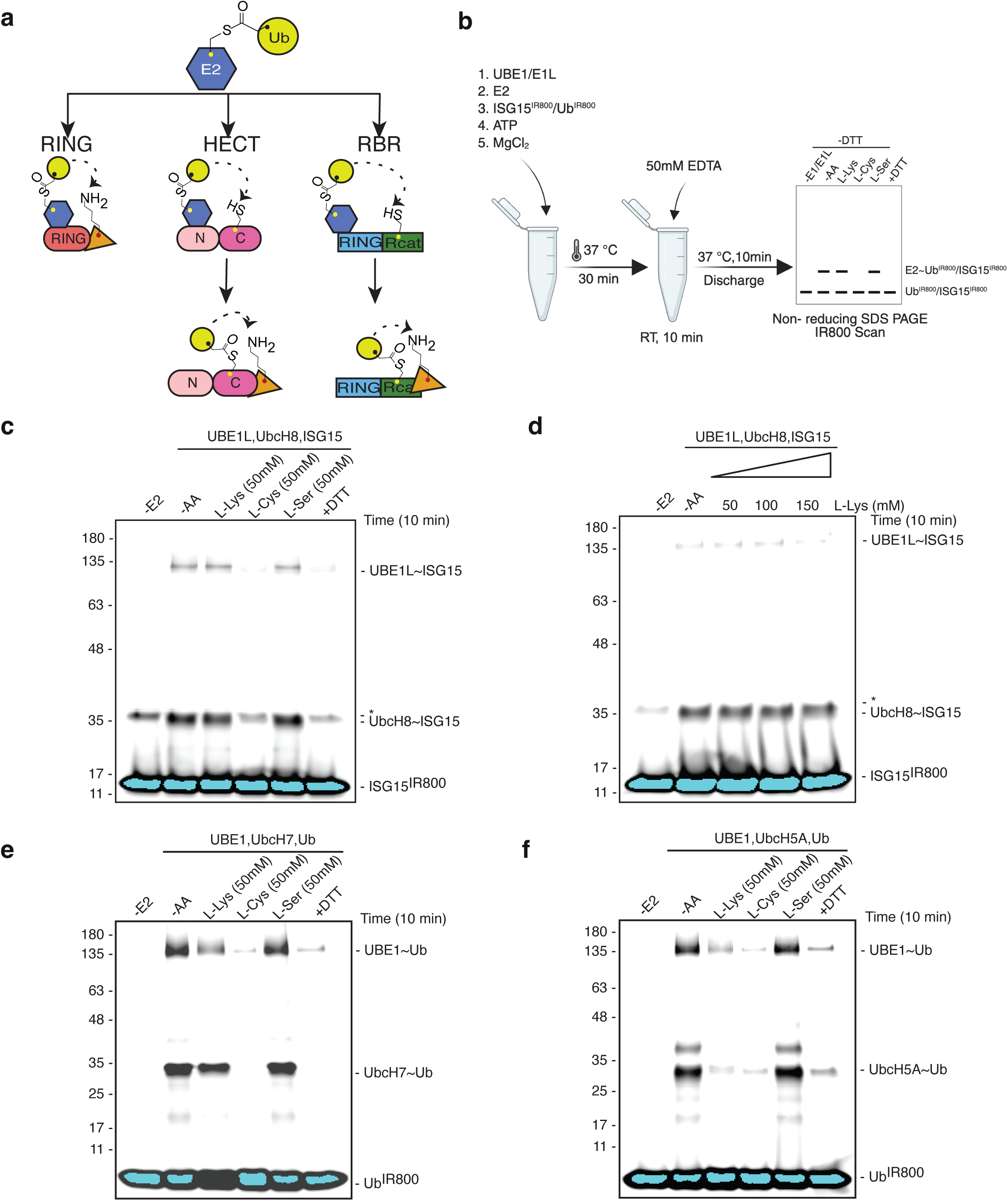
UbcH8∼ISG15 is specifically discharged on cysteine. **a** Schematic illustration of the catalytic mechanisms employed by major E3 ligase families: RING, HECT, and RBR. Acceptor lysine (on E3 or on substrate) is shown in orange. **b** Schematic showing the experimental workflow of the E2 discharge assay using free amino acids. **c** E2 discharge assay using UbcH8∼ISG15 in the presence of various amino acids (50 mM) **d** E2 discharge assay using UbcH8∼ISG15 in the presence of increasing concentrations of L-lys as indicated. **e** E2 discharge assay using UbcH7∼Ub in the presence of various amino acids (50 mM **f** E2 discharge assay using UbcH5A∼Ub in the presence of various amino acids (50 mM)

In this study, we reveal the structural and mechanistic basis for the exclusive specificity of UbcH8∼ISG15 for HECT-family E3 ligases, with HERC5 exhibiting the highest activity, while remaining unreactive toward RING or RBR E3s. In contrast, UbcH8∼Ub preferentially engages RBR E3 ligases (HHARI and Parkin). Our findings demonstrate that a unique closed conformation of UbcH8∼ISG15, stabilized by interactions between the T125 patch of ISG15 (structurally analogous to ubiquitin’s I44 patch) and a divergent crossover helix in UbcH8, enables selective trans-thiolation by HERC5/HERC6/NEDD4 (HECT-family E3s). This conformational distinction explains the incompatibility of ISG15 with RBR-family E3, which specifically recognize ubiquitin’s I44 patch but cannot accommodate ISG15’s T125 patch. We further elucidate how HERC5’s C-lobe achieves exquisite specificity for donor ISG15 recognition during the final E3-to-lysine transfer step, a mechanism that precludes HERC5-mediated ubiquitination activity. These findings provide unprecedented molecular insights into the ISGylation pathway, revealing key evolutionary adaptations that distinguish it from ubiquitination and highlighting potential therapeutic opportunities for modulating ISG15-dependent immune responses.

## Results

### UbcH8∼ISG15 is exclusively reactive with cysteine

Among all E2s, UbcH8 is the only E2 that is compatible with UbE1L to allow formation of ISG15-loaded E2 (UbcH8∼ISG15) ^22–26^. While ubiquitin-conjugating E2 enzymes exhibit broad compatibility with RING, HECT, and RBR E3 ligases^22, 36, 37^, the specificity of ISG15-charged UbcH8 (UbcH8∼ISG15) toward different E3 families remains unclear. Specificity of ubiquitin-conjugating E2s with their respective E3s (HECT/RBR or RING) is also dependent on their ability to react with cysteine/lysine or serine. For example, cysteine-reactive E2s are compatible with HECT or RBR family E3 ligases, whereas lysine-reactive E2s are compatible with RING family E3 ligases. Therefore, if the UbcH8∼ISG15 is compatible with different classes (HECT/RBR and RING) of E3 ligases, UbcH8 must be an E2 that is reactive with both cysteine and lysine. However, UbcH8∼ISG15 discharge assay in the presence of 50 mM L-amino acids (lys/cys/serine) showed discharge of ISG15 with cysteine only, not with lysine or serine (Fig. 1c). Additionally, the use of high concentrations of lysine (up to 150 mM) did not result in the discharge of UbcH8∼ISG15, confirming the specificity of UbcH8 for cysteine (Fig. 1c,d). We also performed discharge assays on UbcH7∼Ub and UbcH5A∼Ub for comparison against UbcH8∼ISG15. Similar to UbcH8∼ISG15, UbcH7∼Ub was specifically reactive with cysteine only, and did not react with lysine or serine (Fig. 1e). However, UbcH5A∼Ub was reactive with both cysteine and lysine (Fig. 1f). The latter is consistent with the fact that UbcH7 is not compatible with RING family E3 ligase, whereas UbcH5A is compatible with RING, HECT, and RBR family E3 ligases (as shown latter). Overall, our data suggest incompatibility of UbcH8 with RING family E3 ligases, which require the E2 to be lysine-reactive.

### ISGylation is catalyzed explicitly by HECT E3 ligase

To check E3 ISG15 ligase selectivity of UbcH8, we used representative active E3 ubiquitin ligases (Extended Data Fig. 2a) from each family (Fig. 2a). Consistent with the discharge assay showing lack of lysine reactivity of UbcH8 (Fig. 1c, d), TRIM25 (RING) did not show any ISGylation activity with UbcH8 (Fig. 2b). Similarly, TRIM25 (RING) did not show ubiquitination activity with UbcH7 (Fig 2c), lacking lysine reactivity (Fig. 1e). Conversely, TRIM25 (RING) showed ubiquitination activity with UbcH5A (Fig 2c), a lysine reactive E2 (Fig. 1f). Despite UbcH8’s ability to react with cysteine (Fig. 1d), active RBR family E3 ligases (HHARI and phospho-Parkin) did not exhibit any significant ISGylation activity (Fig. 2b), while both HHARI and phospho-Parkin showed robust ubiquitination activity with UbcH7 (Extended Data Fig. 2a). Strikingly, HECT-family E3s, HERC5, HERC6, and NEDD4, showed ISGylation activity (Fig. 2b), with HERC5 as the most active ISG15 E3 ligase (Fig. 2b). Also, loss of ISGylation activity upon catalytic cysteine mutation in NEDD4 (Extended Data Fig. 2b) was consistent with the established ubiquitination activity of HECT/RBR family E3 ligases mediated by a catalytic cysteine (Fig. 1a).

**Figure 2.**
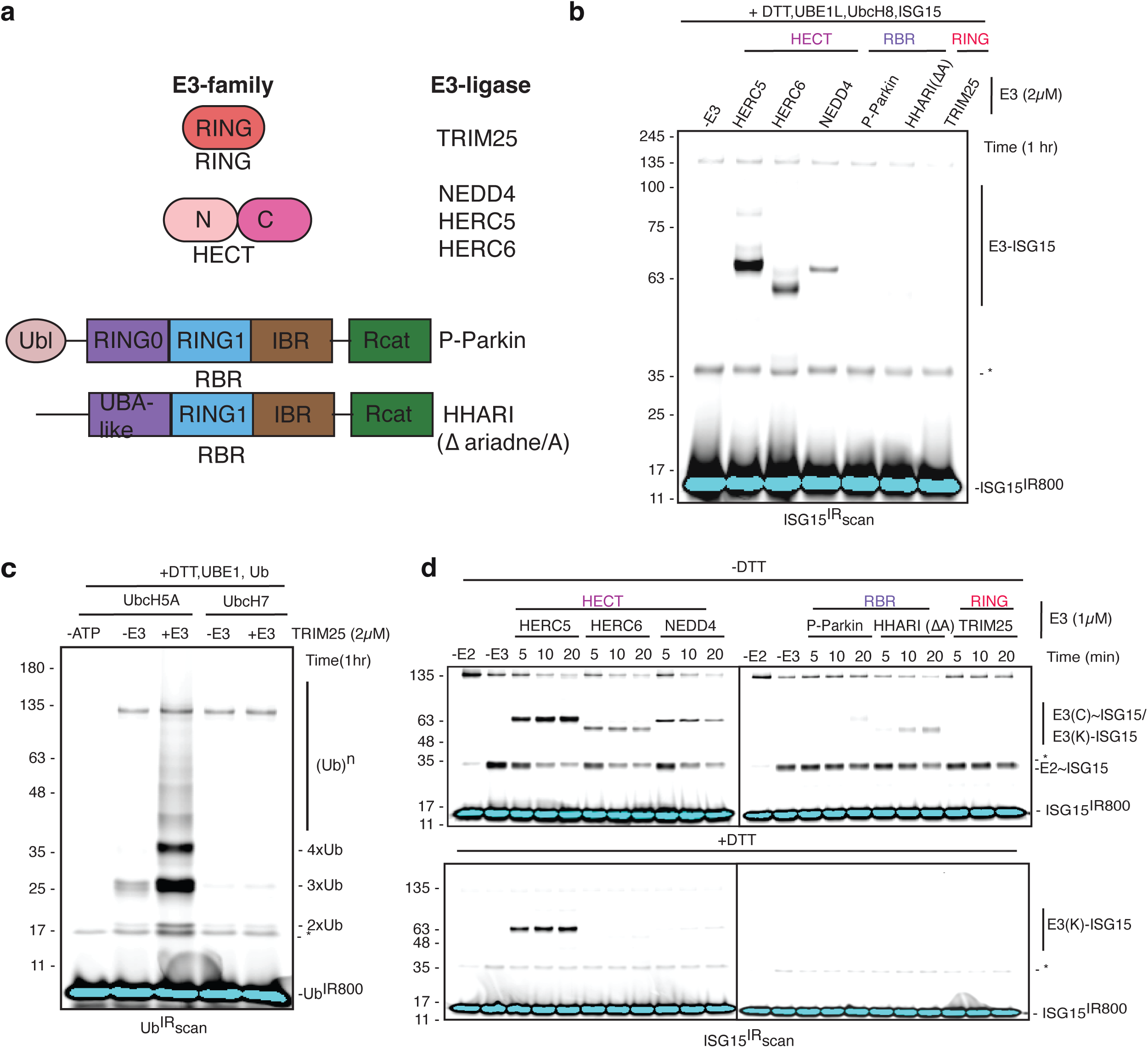
UbcH8-mediated ISGylation is specific to HECT-type E3 ligases. **a** Classification of representative E3 ligases tested for ISGylation activity, categorized by ligase family. **b** ISGylation assays using E3 ligases from different families. IR^800^ scan showing ISGylation activity; an ATP-independent background band is marked with an asterisk (*). **c** Ubiquitination assay using TRIM25 in the presence of various E2s as indicated. IR^800^ scan showing ubiquitination activity; an ATP-independent background band is marked with an asterisk (*). **d** E2 discharge assays using UbcH8∼ISG15 in the presence of various E3 ligases as indicated. Samples were analyzed on non-reducing (-DTT) SDS-PAGE (upper panel) and reducing (+DTT) SDS-PAGE (lower panel). An ATP-independent background band is marked with an asterisk (*).

To further confirm the above findings, we also performed UbcH8∼ISG15 discharge assay in the presence of different family E3 ligases. Consistent with the ISGylation assay (Fig. 2b), UbcH8∼ISG15 was discharged very efficiently by HECT-family E3 ligases (HERC5/HERC6/NEDD4) (Fig. 2d). The addition of DTT in the discharge assay broke the E3(cys)∼ISG15 for HERC6/NEDD4, leaving behind only a fraction of E3(lys)∼ISG15 (Fig. 2d). Conversely, the addition of DTT in the UbcH8∼ISG15 discharge reaction with HERC5 left significantly higher HERC5(lys)-ISG15 (Fig. 2d), suggesting a very fast transfer from cysteine to lysine reaction with HERC5, consistent with ISGylation reaction showing HERC5 as most active E3 ligase (Fig. 2b). In contrast to HECT E3s, RBR family E3s (HHARI/phospho-Parkin) showed extremely poor discharge of UbcH8∼Ub, which was completely vanished in the presence of DTT (Fig. 2d), confirming lack of ISGylation activity (Fig. 2b). Furthermore, consistent with our data in Figure 1 showing lack of lysine reactivity of UbcH8, and Figure 2b showing lack of ISGylation activity with TRIM25, TRIM25 failed to show any significant discharge of UbcH8∼ISG15 (Fig. 1e). These results establish that UbcH8∼ISG15 is selectively recognized by HECT-family E3 ligases, including HERC5/HERC6/NEDD4 (Fig. 2f), and HERC5 as most active E3 ISG15 ligase.

### Change of Ubl (ISG15 to Ub) on UbcH8 changes its specificity from HECT to RBR family E3 ligase

Among all E2 ubiquitin-conjugating enzymes, UbcH8 uniquely partners with UBE1L to form UbcH8∼ISG15 thioester intermediates^21, 31, 32^. Despite a cysteine-reactive E2 (Fig. 1), UbcH8 showed specificity for HECT family E3 ligase, and showed no ISGylation activity with RBR family E3 ligases (Fig. 2). A universal catalytic mechanism of RBR family E3 ligases has been established, where the I44 patch on ubiquitin of E2∼Ub makes critical interactions with RBR family E3 ligase, which is essential for transfer of Ub from E2 to E^37^. We probed whether the differences between Ub and ISG15 are responsible for the HECT specificity observed in Figure 2. UbcH8 demonstrated weaker ubiquitin charging compared to other closest E2s (UbcH6, UbcH7) (Fig. 3a). Notably, RBR-family E3s Parkin/HHARI showed the highest ubiquitination activity with UbcH8 among tested E3s (Fig. 3b-e). However, overall activity was limited by UbcH8’s inefficient ubiquitin charging capacity (Fig. 3a). Interestingly, HECT-family E3 NEDD4 showed robust ubiquitination activity with UbcH6/UbcH7, not with UbcH8 (Fig. 3d), unlike ISGylation activity (Fig. 2). Also, RING-family E3 TRIM25 functioned exclusively with UbcH5A (cysteine and lys reactive), and no ubiquitination was observed with UbcH7/UbcH8 (cys reactive) (Fig. 3e). Together, these findings suggest that UbcH8 exhibits a distinct switch in E3 ligase preference between modification systems, favoring HECT E3s for ISGylation, but displaying its most significant (albeit modest) ubiquitination activity with RBR E3s (Fig. 3f).

**Figure 3.**
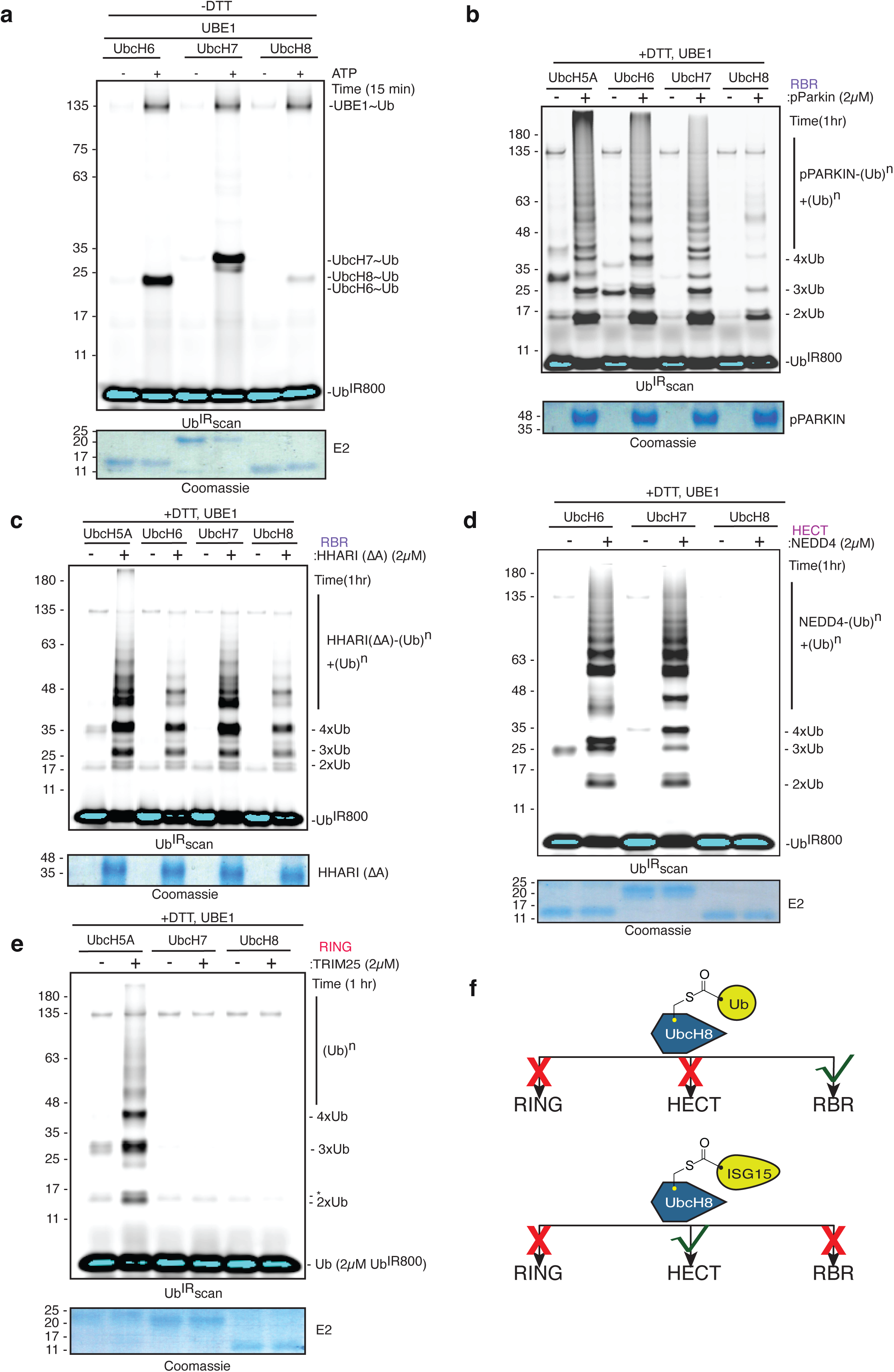
UbcH8 exhibits inverted specificity in ubiquitination reactions. **a** Ubiquitin-charging efficiency of various E2 conjugating enzymes. Top, IR^800^ scan showing formation of E2∼Ub conjugates; bottom, Coomassie-stained gel serves as a loading control. **b–e** In vitro ubiquitination assays using phospho-Parkin (**b**), HHARI (**c**), NEDD4 (**d**), and TRIM25 (**e**) with the indicated E2 enzymes. IR^800^ scan shows ubiquitin conjugation; non-specific bands are marked with an asterisk (*). Coomassie-stained gel showing E2 (lower panels) is used as a loading control. **f** Schematic summary of the key findings from panels **a–e**.

### Crystal structure of the UbcH8∼ISG15 conjugate reveals mechanistic insights into ISGylation

While structures of ubiquitin-charged E2 enzymes have illuminated conformational changes during ubiquitination^33^, the molecular mechanisms governing ISG15 transfer remain undefined. To address this gap, we determined the 2.7 Å crystal structure of UbcH8 covalently conjugated to ISG15 (UbcH8∼cISG15) using a bromopropylamine (3Br) suicide probe of CTD of ISG15 (Extended Data Fig. 3a). The structure of UbcH8∼cISG15 revealed the tail of ISG15 covalently linked with catalytic C86 of UbcH8 (Fig. 4a and Extended Data Fig. 3b). Structure showed extensive interfacial contacts involving UbcH8 residues D28, R54, K64, M67, K69, Q84, and L121, and ISG15 residues Q118, E120, W123, P130, D132, Q134, R153, and L154 (Fig. 4a). Notably, ISG15 adopts a conformation distinct from most E2∼Ub complexes, with its T125 (analogous to ubiquitin’s I44) patch, comprising W123 (analogous to ubiquitin’s R42) and P130 (analogous to ubiquitin’s Q49), buried against UbcH8 (Fig. 4a, b). This contrasts with ubiquitin systems where the I44 patch typically remains surface-exposed in E2∼Ub structures, except in cdc34(E2)∼Ub where it contacts the crossover helix of E2 (Fig. 4b) leading to a closed conformation required for ubiquitination activity. The conformation of UbcH8∼cISG15 observed from crystal structure was similar to the recently reported cryo-EM structure of UBE1L∼UbcH8∼ISG15(t) trans-thiolation intermediate (Fig. 4c). The conformation of free UbcH8∼cISG15/cdc34∼Ub or their complexes with UBE1L/UBE1 showed no major conformational change of ISG15/Ub relative to E2 (Fig. 4 b, c & Extended Data Fig. 3c). The latter suggested that the free state of E2∼ISG15/Ub does not adopt any new conformation on its own. To validate the significance of the conformation of UbcH8∼ISG15 observed from the crystal structure, we mutated residues at the interface of UbcH8 and ISG15. While UbcH8 mutations (D28A/R54A/M67A) had no significant effect on HERC5-mediated ISGylation activity (Fig. 4d), introducing ubiquitin-like residues at ISG15’s T125 patch (W123R/P130Q) severely impaired ISGylation activity (Fig. 4e). We also confirmed that T125 patch mutations did not lead to any significant defect in UbcH8 charging with ISG15 (Extended Data Fig. 3d, e). Overall, our data suggest a critical role of the T125 patch in ISG15 during HERC5 interaction with UbcH8-ISG15, which is required for ISGylation activity of HERC5.

**Figure 4.**
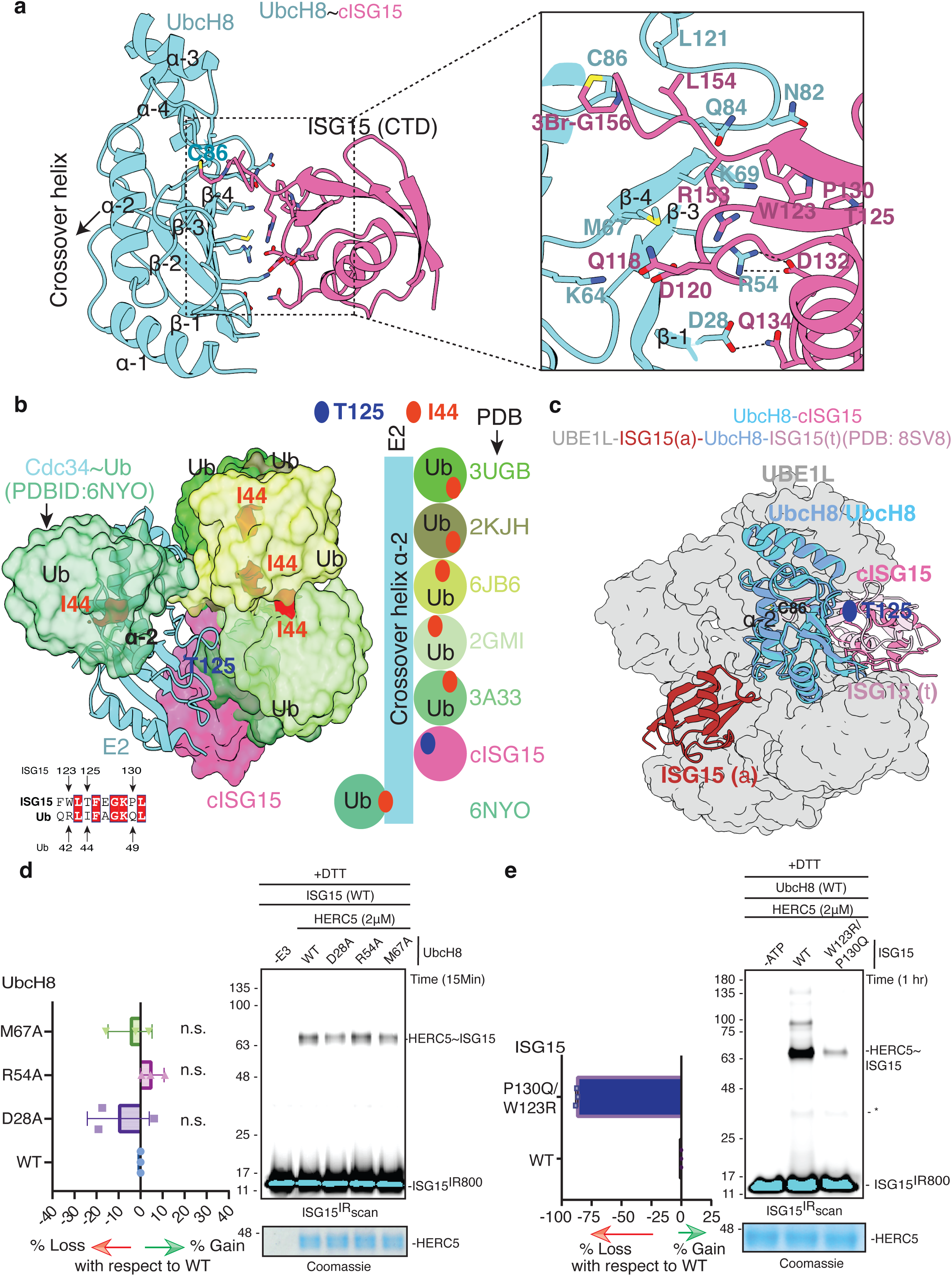
Crystal structure of UbcH8∼cISG15 complex and functional validation. **a** Crystal structure of UbcH8 with the c-terminal domain (CTD) of ISG15 showing key interactions (in the inset) between UbcH8 and ISG15. **b** Overlay of UbcH8(cyan)∼cISG15(pink) structure (this study) with UbcH8∼Ub (PDB 2KJH), UbcH5C∼Ub (PDB 3UGB), Ube2K∼Ub (PDB 6JB6), Ubc13 (PDB 2GMI), UbcH5B∼Ub (PDB 3A33), and cdc34∼Ub (PDB 6NYO). For clarity, only ubiquitin molecules from different E2∼Ub structures are shown. A schematic diagram showing the relative position of the I44/T125 patch of Ub/ISG15 and the crossover helix of E2, from different structures, is also shown. Sequence alignment highlighting divergence in the I44 (Ub)/T125 (ISG15) recognition region. **c** Superposition of the crystal structure of UbcH8∼cISG15(t) on the cryo-EM structure of UBE1L-ISG15(a)-UbcH8-cISG15(t) (PDB 8SV8). Position of T125 is highlighted on the structure of UbcH8∼cISG15/ISG15(t) to show the resemblance in conformations of free or UBE1L-bound states of UbcH8∼cISG15(t). The UBE1L surface (grey) is shown. **d-e** ISGylation activity of HERC5 using UbcH8 (**d**) or ISG15 (**e**) variants as indicated. The left panel shows quantification of HERC5 ISGylation activity (change with respect to WT UbcH8/ISG15). The right panel shows a representative figure of the HERC5 ISGylation assay used for quantification. A non-specific band is indicated (*). The lower panel shows a Coomassie-stained gel as a loading control.

### Closed conformation of UbcH8∼ISG15_donor_ with HERC5 mediates ISG15 E2-E3 trans-thiolation

Ubiquitin-charged E2 enzymes adopt distinct conformational states upon E3 ligase engagement, which in turn govern enzyme specificity^39–44^. RING-family E3s stabilize a closed E2∼Ub conformation with ubiquitin’s I44 patch contacting the E2 crossover helix, while RBR-family E3s maintain an open state with I44 bound to the E3 catalytic domain (Extended Data Fig. 1a, b). Crystal structures of HECT family E3 with E2∼Ub revealed a distinct conformation of ubiquitin with surface exposed I44 patch (Extended Data Fig. 1c). However, recent cryo-EM structure of HECT (pub2) with E2(ubc4/UbcH5)∼Ub^39^ revealed a closed E2∼Ub conformation similar to that of RING family E3 ligase (Fig. 5a, Extended Data Fig. 1a). To investigate whether a similar mechanism governs ISGylation, we modeled the HERC5 and UbcH8∼ISG15 complex by superposition on the cryo-EM structure of the HECT (pub2) and E2(ubc4)∼Ub complex (Fig. 5b).

**Figure 5.**
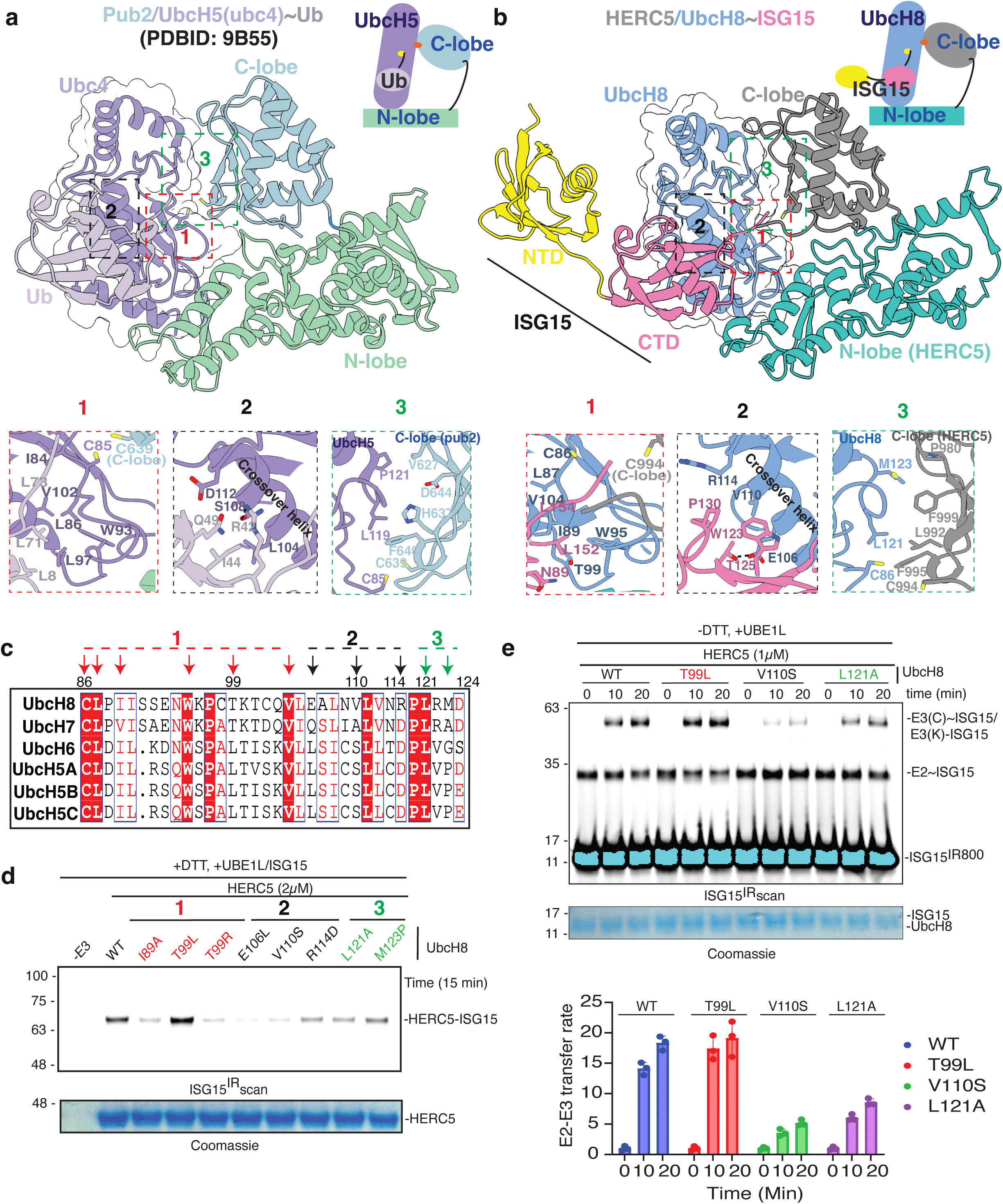
A closed conformation of UbcH8∼ISG15 determines the specificity between UbcH8 and HECT. **a** Cryo-EM structure of the closed conformation of UbcH5∼Ub bound to Pub2 (HECT) (PDB 9B55). **b** Model of UbcH8∼ISG15 and HERC5 complex showing analogous interface architecture. The major sites between E2 and Ub/ISG15 or E3 are boxed as Site 1 (red), Site 2 (black), and Site 3 (green), respectively (**a, b**). The schematic diagram on top depicts the conformation of the structures (**a, b**). The catalytic cysteines of E2 and E3 are highlighted as yellow and orange shapes, respectively, in the schematic above. **c** Sequence alignment of UbcH8 with other ubiquitin-conjugating E2 enzymes. Residues contributing to interaction sites 1, 2, and 3 (defined in **a** and **b**) are indicated by arrows colored red, black, and green, respectively. **d** ISGylation activity of HERC5 with wild-type and mutant UbcH8 variants targeting key interface residues from all three sites. **e** E2 discharge assay monitoring the rate of ISG15 transfer from UbcH8 to HERC5. Wild-type and mutant forms of UbcH8, targeting Site 1 (T99L), Site 2 (V110S), and Site 3 (L121A), were analyzed over a time course as indicated (upper panel). Quantification represents the mean of three independent experiments (lower panel), fold change was calculated relative to the 0-minute time point.

Comparative structural analysis identified three conserved interaction sites as Site 1, Site 2, and Site 3 (Fig. 5a, b). Site 1 was characterized by conserved hydrophobic interactions, L8/L71/L73 of Ub with L97/W93/L86/V102/I84 of UbcH5 (Fig. 5a), and N89/L152/L154 of ISG15 with T99/W95/I89/V104/L87 of UbcH8 (Fig. 5b). Site 2 showed contrasting differences between ubiquitin or ISG15 and the crossover helix of their respective E2s to accommodate the donor state of E2∼Ub/ISG15_(donor)_ (Fig. 5a, b). Site 2 highlighted the structural divergence between ISG15 and Ub in how they engage the E2 crossover helix in their respective donor states. Specifically, Ub residues I44, R42, and Q49 interact with L104, S108, and D112 of UbcH5, whereas ISG15 residues T125, W123, and P130 contact E106 and V110 of UbcH8 (Fig. 5a, b). Site 3 defined the catalytic interface, where interactions between the E2 and the E3 C-lobe position their respective catalytic cysteines for trans-thiolation (Fig. 5a, b). In the ubiquitin system, Site 3 included L119 and P121 of UbcH5 contacting F540, H637, D644, and V627 of the E3 Pub2 (Fig. 5a). In the ISG15 system, Site 3 included L121 and M123 of UbcH8 contacting F995, L992, F999, and P980 of HERC5 (Fig. 5b).

Sequence alignment of UbcH8 with other ubiquitin-conjugating enzymes showed that Site 1 is highly conserved across E2s. In contrast, Site 2 shows distinct adaptations in UbcH8 that accommodate ISG15’s unique surface on the T125 patch (Fig. 5c). Site 3 shows conservation of L121 (Fig. 5c) among E2s, however, residues corresponding to M123 of UbcH8 are mutated to less bulky side chains in other E2s (Fig. 5c). Functional mutagenesis confirmed the importance of all three sites on UbcH8. Mutations (I98A/T99R) in Site 1 perturbing the conserved hydrophobic core resulted in loss of ISGylation activity (Fig. 5d). In contrast, the T99L mutation (mimicking ubiquitin conjugating enzymes) of UbcH8 enhanced HERC5-mediated ISGylation activity by favoring interactions in the hydrophobic pocket (Fig. 5d, 5a-c, Extended Data Fig. 4a). The latter also confirmed that T99 on UbcH8 is a naturally attenuating variant. Mutations (E106L/V110S/R114D), mimicking ubiquitin conjugating enzymes, in the Site 2 involving crossover of UbcH8 drastically perturbed HERC5-mediated ISGylation activity (Fig. 5d, Extended Data Fig. 4a). Similarly, mutations (L121A/M123P) of UbcH8 in Site 3 perturbed HERC5-mediated ISGylation activity, supporting the critical role of this interface in catalysis (Fig. 5d). Additionally, UbcH8 mutations (sites 1/2/3) showed no significant defects on UbcH8 charging with ISG15 (Extended Data Fig. 4b), confirming intact E1-E2 trans-thiolation.

We also probed the role of the above interactions on the transfer rate of ISG15 from E2 to E3 under non-reducing conditions (Fig. 5e). V110S on the crossover helix in Site 2, perturbing the close conformation of UbcH8∼ISG15, severely impaired transfer of ISG15 from E2 to E3 leading to accumulation of UbcH8∼ISG15 and loss of HERC5(cys)∼ISG15/ HERC5(lys)-ISG15 (Fig. 5e). Similarly, L121A, which perturbed the positioning of catalytic cysteines of E2 and E3 in Site 3 (Fig. 5a, b), markedly reduced transfer of ISG15 from E2 to E3 (Fig. 5e). Conversely, T99L mutation on UbcH8 enhanced the rate of ISG15 transfer from E2 to E3 (Fig. 5e). Results of UbcH8∼ISG15 discharge assay (Fig. 5e) were consistent with our observations in Figure 5d. These results precisely map the structural determinants of ISGylation specificity, demonstrating how ISG15’s unique surface chemistry and UbcH8’s specialized crossover helix coordinate to enforce productive E2-E3 trans-thiolation during ISGylation reaction.

### Selective ISG15 conjugation is governed by a specialized donor-binding pocket in HERC5

After understanding the mechanism of ISG15 transfer from UbcH8 to HERC5, we next probed whether, similar to other HECT family E3 ligases, HERC5 could also work as a ubiquitin E3 ligase. While HERC5 showed a very robust ISGylation activity with UbcH8 (Fig. 2), it failed to support HERC5-mediated ubiquitination (Extended Data Fig. 5a). Furthermore, HERC5 did not show any significant ubiquitin ligase activity regardless of E2 partners (UbcH5A, UbcH6, UbcH7) (Extended Data Fig. 5a). We wanted to understand the mechanism of HERC5-mediated specificity for ISGylation. To understand this, we modeled the structure of ISG15-loaded HERC5 using Alfafold3 and compared it with the crystal structure of ubiquitin-loaded NEDD4 (Fig. 6a, b). Interactions between ISG15 and HERC5 were mediated between the C-lobe of HERC5 and the CTD of ISG15, without involving the NTD of ISG15 (Fig. 6a). Comparative structural analysis of ISG15-bound HERC5 and ubiquitin-bound NEDD4 revealed conserved overall architecture but critical differences at the E3 and the donor ISG15/Ub interface (Fig. 6a, b). The HERC5∼ISG15 interaction core involved ISG15 residues K90/N89 and HERC5 residues E1015/P988/D987, distinct from the NEDD4∼Ub interface involving I36/D39 of Ub and M887/L860/K859 of NEDD4 (Fig. 6a, b). Sequence alignment confirmed HERC5’s uniqueness with P988/E1015 among HECT E3s (Fig. 6c), which could contribute towards HERC5 specificity against ISG15.

**Figure 6.**
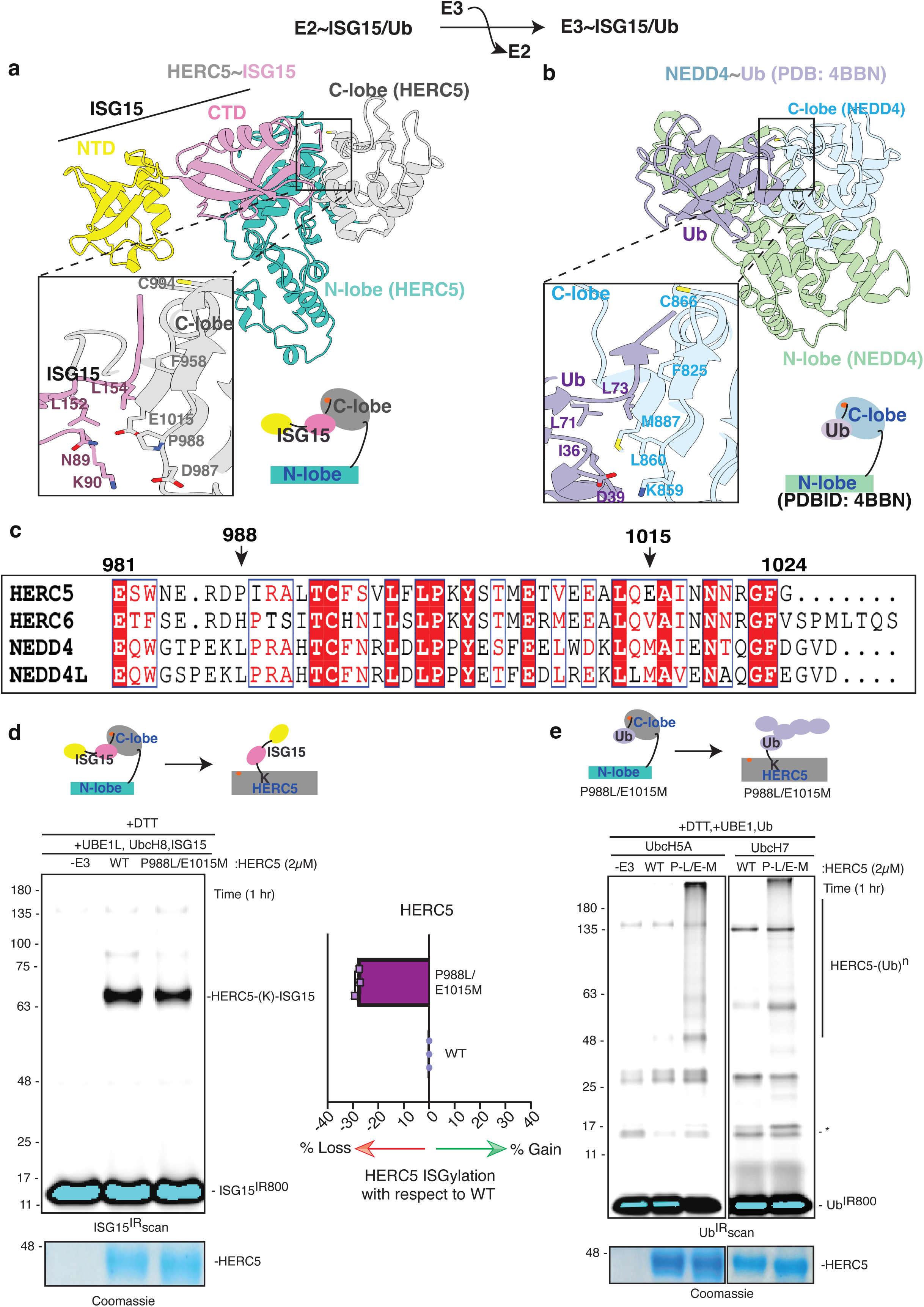
A distinct donor ISG15-binding site in the C-lobe of HERC5 governs ISGylation specificity. **a** Predicted structure of the HERC5∼ISG15 complex generated using AlphaFold3. The N- and C-terminal domains (NTD and CTD) of ISG15 are shown in distinct colors. The model positions the ISG15 C-terminal tail near the catalytic cysteine (C994) of HERC5. An inset highlights key interactions between ISG15 and HERC5. A schematic of the complex is also shown. **b** Crystal structure of the NEDD4∼Ub complex (PDB 4BBN), illustrating the Ub C-terminal tail engaged with the catalytic cysteine (C866) of NEDD4. The inset shows detailed interactions within the interface. A schematic representation accompanies the structure. **c** Sequence alignment of the HERC5 C-lobe with corresponding regions of representative HECT E3 ligases. Residues implicated in donor (ISG15 or Ub) recognition are indicated with arrows. **d** ISGylation assays with HERC5 variants validate the structural model. Left, a representative gel showing ISGylation activity; Coomassie-stained gel below serves as a loading control. Right, quantification of ISGylation activity from three biological replicates. A schematic of the reaction captured from the assay is shown above. **e** Ubiquitination assays using HERC5 variants to test specificity inferred from structural analysis. A schematic of the reaction captured from the assay is shown above.

To confirm the structural findings, P988L/E1015M mutation, a NEDD4-mimic, on HERC5 resulted in loss of HERC5 ISGylation activity (Fig. 6d). Similarly, L1281P/M1370E mutation, a HERC5-mimc, on NEDD4 ablated ubiquitination activity of NEDD4 (Extended Data Fig. 5b). Furthermore, while WT HERC5 did not show any significant ubiquitination, a robust ubiquitination by P988L/E1015M HERC5 with both UbcH5A and UbcH7 (Fig. 6e). The latter observation confirmed the role of the C-lobe in making HERC5 a specific E3 ISG15 ligase (Fig. 6e). Furthermore, unlike UbcH8 mutations perturbing E2-E3 trans-thiolation leading to accumulation of UbcH8∼ISG15 in Fig. 5e, P988L/E1015M HERC5 did not show any significant defect on discharge of UbcH8∼ISG15 (Extended Data Fig. 5c). However, similar to other HECT-family E3 ligases (Fig. 2d), P988L/E1015M HERC5 resulted in loss of E3(lys)-ISG15 (Extended Data Fig. 5c), pinpointing the role of above interactions in the final lysine conjugation step of ISGylation reaction (Fig. 1a). These results establish that HERC5’s specialized C-lobe interface enforces ISG15 specificity during the terminal conjugation step, explaining its exclusive ISGylation activity among E3 ligases.

### Conserved E2 recognition by the HECT N-lobe enables ISG15 transfer by non-cognate HECT E3 ligases

Ubiquitin-conjugating enzymes (E2s) exhibit high sequence and structural similarity (Extended Data Fig. 6), enabling broad compatibility with various E3 ubiquitin ligases. For instance, NEDD4 (HECT-type E3) efficiently ubiquitinates with UbcH6 or UbcH7 but shows no activity with UbcH8, unlike phospho-Parkin or HHARI (Fig. 3b-e). Conversely, NEDD4 discharges UbcH8∼ISG15 to mediate ISGylation, although weaker than HERC5 (Fig. 2b). This prompted us to investigate how NEDD4 is compatible with different E2s for different donor molecules. To explore this, we used AlphaFold3 to model UbcH6/NEDD4 and UbcH8/HERC5 complexes. Both structures revealed striking similarities, with interactions primarily involving the HECT domain’s N-lobe and the E2 (Fig. 7a,b). Key residues in the UbcH6/NEDD4 interface included UbcH6’s F108/K109 and NEDD4’s L1115/I1117/I1107/L1104/Y1100/E1099 (Fig. 7a). Similarly, the UbcH8/HERC5 interface relied on UbcH8’s F116/K117 and HERC5’s I834/F828/L820/L817/L813/D812 (Fig. 7b). These interaction sites were greatly conserved across E2s (Extended Data Fig. 6) and HECT E3s (Fig. 7c), explaining their promiscuity. ITC measurements further supported this, showing comparable binding affinities between NEDD4 and UbcH6/UbcH7/UbcH8 (Fig. 7d). Since HECT E3s and E2s share a conserved binding mechanism, and ISGylation closely resembles ubiquitination in requiring a closed E2∼Ub/ISG15_donor_ conformation for E2-E3(HECT) trans-thiolation (Fig. 5), NEDD4 can mediate both reactions when paired with appropriate E2-donor (Ub/ISG15) combinations (Figs. 2 & 3). However, specificity is ultimately determined by distinct conformational changes required for donor (Ub/ISG15) transfer from E3 to lysines (Fig. 6).

**Figure 7.**
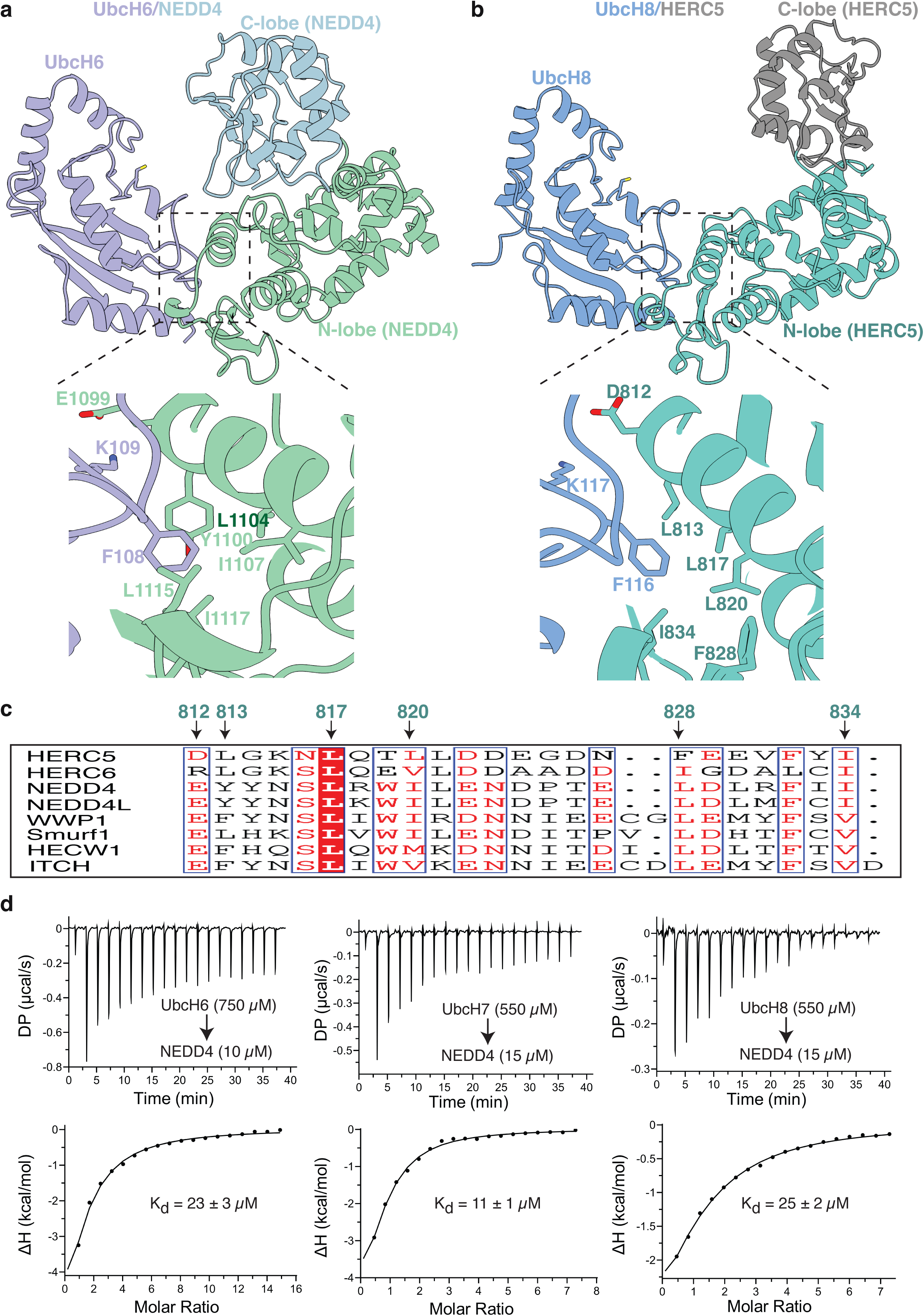
Conserved interfaces mediate E2–HECT E3 interactions. **a** AlphaFold3 structure of the UbcH6/NEDD4 complex generated using AlphaFold3, highlighting key interactions at the E2–E3 interface. **b** AlphaFold3 structure of the UbcH8/HERC5 complex, showing conserved contacts between the E2 enzyme and the HECT E3 ligase. **c** Sequence alignment of the N-lobe regions from representative HECT family E3 ligases. Residues contributing to the E2–E3 interface, as identified in panels **a** and **b**, are indicated with arrows. **d** Isothermal titration calorimetry (ITC) analysis of NEDD4 binding to UbcH6 (left), UbcH7 (middle), and UbcH8 (right). Protein concentrations used in the sample cell and syringe are indicated in parentheses.

## Discussion

ISGylation is a critical post-translational modification that plays a significant role in host defense against viral infections and cancer^1–6^. Despite its importance, the molecular mechanisms underlying ISGylation remain poorly understood. Previous studies have demonstrated that among various E2 ubiquitin-conjugating enzymes, only UbcH8 is charged by UBE1L^21^. However, the specificity between UbcH8 and E3 ligases (which also function as ubiquitin ligases) remains unclear.

Here, we found that UbcH8 was compatible with multiple HECT-family E3 ligases (HERC5, NEDD4, and HERC6). In contrast, other proposed ISG15 E3 ligase TRIM25 (RING), or RBR-family E3 ligase (HHARI/phospho-Parkin) exhibited no ISGylation activity (Fig. 2). UbcH8 exhibited a shift in specificity between ISGylation and ubiquitination when tested with different E3 ligases (Figs. 2 & 3). Our findings revealed that when UbcH8 is loaded with ISG15, it adopts a closed conformation upon binding with the HECT-family E3 ligase, similar to the closed conformation of E2∼Ub (Fig. 5). The closed conformation is crucial for E2-E3 trans-thiolation in both ubiquitination/ISGylation mediated by HECT family E3 ligases. This closed conformation of UbcH8 is facilitated by the interaction between the crossover helix of UbcH8 and the T125 patch of ISG15 (Fig. 5). The T125 patch of ISG15 corresponds to the I44 patch on ubiquitin, and variations in the crossover helix of UbcH8 make it specific for ISG15 (Fig. 5), while similar affinities between HECT and E2s make it promiscuous with HECT family E3 ligase (Fig. 7). Consequently, while UbcH8 demonstrated ISGylation activity with multiple HECT E3 ligases (Fig. 2), it did not exhibit ubiquitination with HECT E3 ligases (Fig. 3 & Extended Data Fig. 5a). This also accounts for UbcH8∼Ub compatibility with RBR-family (HHARI/phospho-Parkin) mediated ubiquitination activity, as E2∼Ub with RBR family E3 ligases are known to adopt an open conformation facilitated by interactions between the I44 patch on ubiquitin and the E3 ligase (Extended Data Fig. 1b,37).

Despite the necessity of a closed conformation for both RING and HECT E3-mediated reactions, UbcH8 fails to support ISGylation via TRIM25 (Fig. 2). This incompatibility stems from UbcH8’s exclusive reactivity toward cysteine (Fig. 1c,d), whereas RING-type E3 ligases require lysine-reactive E2s that can discharge ubiquitin directly onto substrate lysines (Figs. 1f, 2c, and ^33^). These findings reveal an additional layer of specificity in the ISGylation cascade, defined not only by the UBE1L–UbcH8 pairing but also by the interplay between the donor (Ub or ISG15) on UbcH8 (E2) and E3 conformation preferences (Fig. 8).

**Figure 8.**
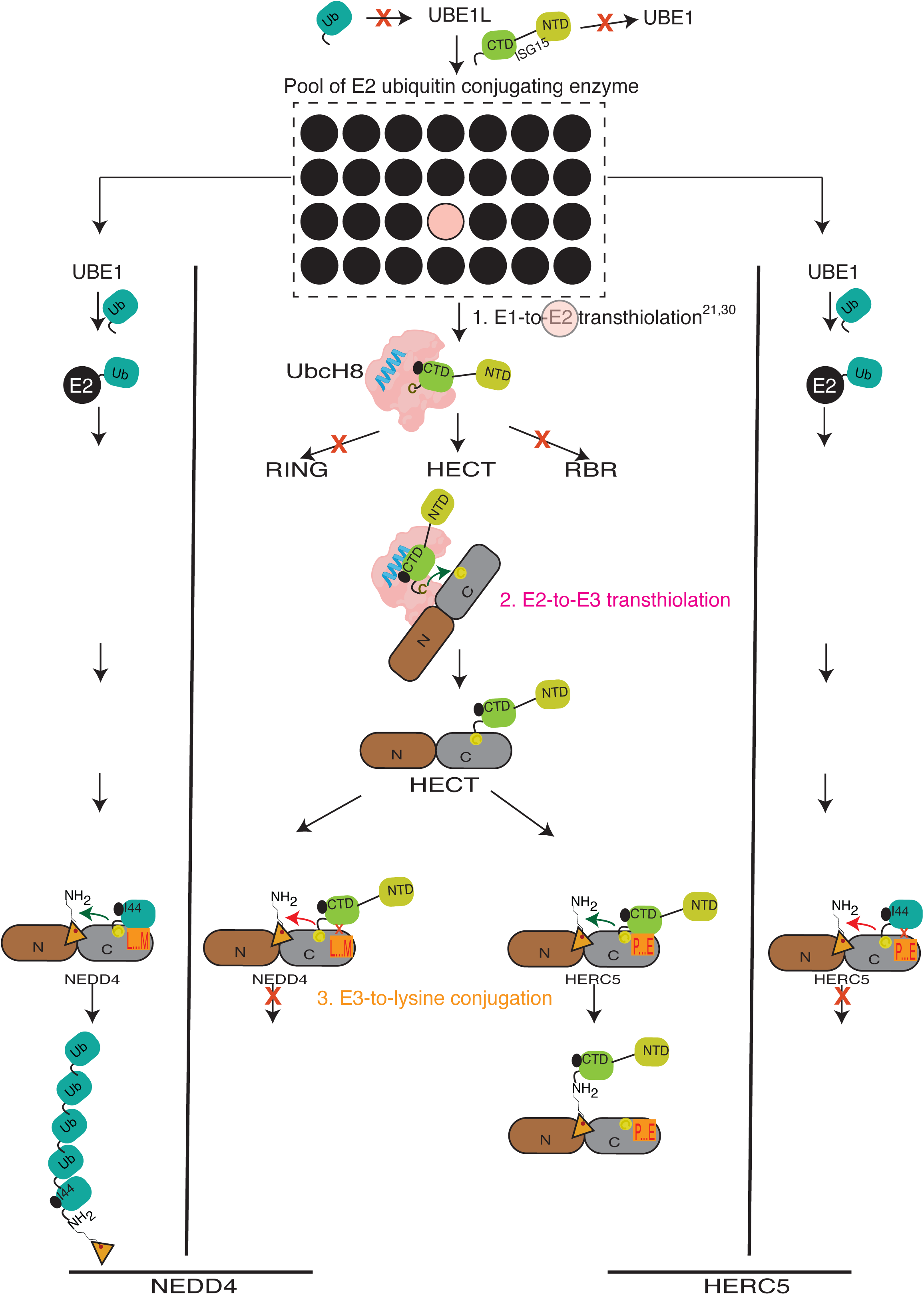
Proposed model illustrating hierarchical specificity in the ISGylation cascade. Schematic representation of the ISGylation pathway highlighting distinct levels of specificity, including E1–E2, E2–E3, and donor (ISG15) recognition on E3. The model integrates structural and biochemical findings to depict how specificity is achieved at multiple steps in the enzymatic cascade. Crossover helix (blue) on UbcH8 (pink) is shown. Selectivity/specificity is indicated as (X).

Among tested E3 ligases, HERC5 exhibited the most robust ISGylation activity (Fig. 2) but lacked ubiquitination activity with different E2 conjugating enzymes, including UbcH8 (Fig. 6e & Extended Data Fig. 5a), establishing it as a highly selective ISG15 E3 ligase. Mechanistically, after E2-E3 trans-thiolation, ISG15 is stabilized on HERC5’s C-lobe, which is distinct from other E3 ligases (Fig. 6). This distinct site on HERC5 preferentially accommodates ISG15 over ubiquitin. Notably, mutating key residues to enhance ubiquitin binding on the C-lobe transformed HERC5 into an active ubiquitin ligase (Fig. 6e). This interaction is essential for the final transfer step in ISGylation/ubiquitination reaction of lysine conjugation (Fig. 8). Collectively, our data reveal an E3-dependent specificity mechanism governing ISGylation (Fig. 8).

In summary, our study elucidates the molecular mechanisms governing ISGylation, uncovering key specificity determinants (Fig. 8) that could help the development of targeted therapeutic interventions.

## Methods

### Molecular Biology

For expression in insect cells, full-length human UBE1L and the HECT domain of human HERC5 were amplified from HEK293T cDNA and subcloned into the pFastBac HT B vector (Invitrogen), which includes an N-terminal 6×His tag followed by a TEV protease cleavage site. Genes encoding UbcH8, UbcH7, UbcH6, HERC6, NEDD4, TRIM25, ISG15, ubiquitin, and Parkin were amplified from human cDNA and inserted into the pET15b or pET22 vector (Novagen) for bacterial expression. Site-directed mutagenesis was used to generate all point mutants. To enable site-specific fluorescent labeling, an N-terminal cysteine was introduced into ISG15 and ubiquitin by mutagenesis. For synthesis of the cISG15-3Br suicide probe, the C-terminal domain of ISG15 (residues 80–156) was cloned into the pTXB1 vector (New England Biolabs). The following plasmids were obtained from Addgene: UBE1 (Addgene plasmid #34965; gift from Cynthia Wolberger), UbcH5A (Addgene plasmid #15782; gift from Wade Harper), and NEDD4L (Addgene plasmid #165103; gift from Arthur Haas). Domain boundaries of proteins used in this study are provided in the Extended Data Table 2.

### Protein expression and purification

Recombinant UBE1L and HERC5 were expressed using the Baculovirus Expression Vector System (BEVS, Thermo Fisher Scientific). Baculoviruses were generated in ExpiSf9 cells (a gift from Dr. Debasish Nayak) and used to infect ExpiSf9 cultures grown in ExpiCD medium (Gibco, A3767803) at a density of 5–10 × 10⁶ cells/mL. Cultures were incubated at 27 °C for 4–5 days before harvesting. Cell pellets were stored at –80 °C until lysis. Cells were lysed by sonication in buffer containing 50 mM Tris (pH 7.5), 300 mM NaCl, 5% glycerol, 5 mM β-mercaptoethanol (β-ME), 1 mM AEBSF, DNase I, and 10 μg/ml leupeptin. Lysates were clarified by centrifugation at 18,000 × g for 40 min at 4 °C. Supernatants were incubated with Ni-NTA resin for 2 h at 4 °C. Proteins were eluted using 300 mM imidazole in 50 mM Tris (pH 7.5), 150 mM NaCl, and 5 mM β-ME. Eluted proteins were buffer-exchanged into 20 mM Tris (pH 8.0), 200 mM NaCl, and 2 mM β-ME, flash-frozen in liquid nitrogen, and stored at – 80 °C. Other recombinant proteins were expressed in *E. coli* BL21(DE3) cells. Cultures were grown at 37 °C to an OD₆₀₀ of 0.6–0.8, induced with 0.3 mM IPTG, and incubated overnight at 18 °C. Cells were harvested by centrifugation (5,000 × g, 15 min, 4 °C), lysed by sonication, and clarified by centrifugation. Proteins were purified using Ni-NTA or glutathione-Sepharose (GSH) affinity chromatography, followed by overnight cleavage of affinity tags at 4 °C using SENP1 or PreScission protease. Further purification was performed by ion-exchange and/or size-exclusion chromatography using HiLoad 16/600 Superdex 75 pg or Superdex 200 pg columns (Cytiva) in 20 mM Tris (pH 8.0), 100 mM NaCl, and 2 mM β-ME. Purified proteins were flash-frozen and stored at –80 °C. Purification protocols for phospho-Parkin, UbcH7, and ubiquitin were as described previously^40, 41^.

### Preparation of IR800-labeled ISG15 and ubiquitin

For use in ISGylation and ubiquitination assays, ISG15 and ubiquitin were fluorescently labeled with IR800 dye as previously described^40, 41^. A cysteine residue was engineered at the N-terminus of each protein to enable site-specific labeling. The native cysteine (Cys78) in human ISG15 was mutated to serine to prevent non-specific conjugation. Proteins were buffer-exchanged using a HiLoad 16/600 Superdex 75 pg column (Cytiva) equilibrated in PBS containing 1 mM TCEP. Labeling was carried out by incubating proteins with sulfo-Cyanine7.5 maleimide (Lumiprobe) for 30 min at room temperature under an argon atmosphere, following the manufacturer’s protocol^42^. Labeled proteins were dialyzed overnight at 4 °C and further purified by SEC using a Superdex Increase 75 10/300 GL column (Cytiva) equilibrated in 25 mM Tris (pH 7.5), 75 mM NaCl. Purified proteins were concentrated, flash-frozen in liquid nitrogen, and stored at –80 °C.

### E2 charging assay

E2∼ISG15/Ub conjugates were generated in a 20 μL reaction containing 0.25 μM E1 enzyme (UBE1 for ubiquitin, UBE1L for ISG15), 1 μM E2 enzyme, and 2 μM fluorescently labeled ubiquitin or ISG15. Reactions were carried out in charging buffer composed of 20 mM HEPES (pH 7.5), 50 mM NaCl, 5 mM MgCl₂, and 5 mM ATP. The E1 enzyme was first pre-activated by incubation at 25 °C for 5 min, followed by the addition of E2 and further incubation for 10 min at 25 °C. Reactions were terminated by adding non-reducing SDS-PAGE sample buffer. Samples were resolved by SDS-PAGE on a 15% gel and visualized using a LI-COR Odyssey Infrared Imaging System. Band intensities were quantified using ImageJ software.

### E2 discharge assay

Assays showing Ub or ISG15 discharge from E2 were performed in 50 mM Tris (pH 7.5) supplemented with 5 mM ATP and 5 mM MgCl₂. For charging, 1 μM E2 was incubated with 2 μM fluorescently labeled ubiquitin or ISG15 and 0.25 μM UBE1 (for Ub) or UBE1L (for ISG15) at 30 °C for 15 min. Charging was quenched by addition of 50 mM EDTA, followed by 10 min incubation at 25 °C. For discharge, E3 ligases were added at a final concentration of 1 μM, and reactions were incubated at 30 °C. Similarly, 50mM (or as indicated) L-cys/lys/ser were added to E2∼ISG15/Ub to initiate the discharge of E2, thereby capturing E2 reactivity with the respective amino acids. Time-point samples were taken as indicated, and reactions were stopped by adding non-reducing SDS-PAGE loading buffer. Samples were resolved on a 6–15% SDS-PAGE gradient gel and analyzed using LI-COR Odyssey Infrared Imaging System.

### ISGylation or Ubiquitination Assay

ISGylation and ubiquitination assays were performed in a 20 μL reaction volume containing 0.25 μM E1 enzyme (UBE1L for ISG15 or UBE1 for ubiquitin), 1 μM E2 conjugating enzyme, 2 μM E3 ligase, and 2 μM infrared dye-labeled ISG15^IR800^ or Ub^IR800^. Reactions were carried out at 30 °C in assay buffer composed of 25 mM Tris (pH 7.5), 50 mM NaCl, 10 mM MgCl₂, 0.5 mM DTT, and 5 mM ATP. Reactions were stopped by the addition of SDS-PAGE loading buffer supplemented with 20 mM DTT, followed by denaturation at 95 °C for 5 min. Proteins were resolved on 6–15% SDS-PAGE gradient gels and visualized using the LI-COR Odyssey Infrared Imaging System.

### Isothermal Titration Calorimetry (ITC)

Protein–ligand binding affinities were measured using a MicroCal PEAQ-ITC instrument (Malvern Panalytical) at 25 °C. All proteins were dialyzed extensively into phosphate-buffered saline (PBS, pH 7.4) supplemented with 2% (v/v) glycerol and 2 mM β-mercaptoethanol. Titrations were performed by injecting 23 successive 2 μL aliquots of ligand into the calorimetric cell containing the binding partner, with 120-second intervals between injections. All experiments were conducted with constant stirring at 750 rpm. Raw data were analyzed using the Microcal PEAQ-ITC analysis software and fitted to a single-site binding model to derive thermodynamic parameters and the equilibrium dissociation constant (Kd).

### Synthesis and purification of cISG15-3Br

The synthesis and purification of cISG15-3Br were carried out as previously described^40, 43^, with minor modifications. The C-terminal domain of ISG15 (residues 80–156; referred to as cISG15) was cloned into the pTXB1 vector as a fusion with the Mxe-intein–chitin binding domain (CBD) and a C-terminal His tag. The resulting construct, cISG15(80–156)–intein– CBD–His, was expressed in *Escherichia coli* BL21(DE3) cells. Protein expression was induced at an optical density (OD₆₀₀) of 0.7 by addition of 0.2 mM IPTG, followed by incubation at 16 °C for 16 h. Cells were lysed in buffer containing 50 mM HEPES (pH 6.5) and 200 mM NaCl. The fusion protein was purified using Ni-NTA affinity chromatography, and cleavage was induced by overnight incubation of the resin with elution buffer containing 50 mM HEPES (pH 6.5), 200 mM NaCl, and 0.15 M sodium 2-mercaptoethanesulfonate (MESNa). The eluted thioester intermediate was subsequently reacted with 3-bromopropylamine hydrobromide (Sigma-Aldrich) at 25 °C for 3 h at pH 8.0 to generate cISG15-3Br. The final product was purified by size-exclusion chromatography using a HiLoad 16/600 Superdex 75 pg column (Cytiva) pre-equilibrated in 50 mM HEPES (pH 6.5) and 200 mM NaCl. Fractions containing cISG15-3Br were pooled, concentrated, flash-frozen in liquid nitrogen, and stored at –80 °C until use.

### Preparation and Crystallization of UbcH8∼cISG15 Complex

The UbcH8∼ISG15 conjugate was generated using a UbcH8 variant (C97S/C101S), retaining only the catalytic cysteine (C86). The complex was formed by incubating UbcH8 with a two-fold molar excess of cISG15-3Br at 25 °C for 4 hours. The covalent complex was purified by size-exclusion chromatography (HiLoad 16/600 Superdex 75 pg, Cytiva) in storage buffer (25 mM Tris-HCl pH 7.5, 75 mM NaCl). For crystallization trials, the purified complex (9.5 mg/mL) was screened using sitting-drop vapor diffusion at 18 °C. Initial crystal hits were obtained in 0.1 M MES (pH 6.5), 15% PEG 550 MME. Diffraction-quality crystals were subsequently optimized using microseeding in hanging-drop format. Prior to X-ray data collection, crystals were cryoprotected in mother liquor supplemented with 25% glycerol and flash-cooled in liquid nitrogen.

### Data processing and Model building

Diffraction data were collected at 100 K using beamline ID30B at the European Synchrotron Radiation Facility (ESRF, Grenoble). Raw diffraction images were processed with XDS^44^ and scaled using Aimless^45^ from the CCP4 suite^46^. The UbcH8∼cISG15 structure was solved by molecular replacement in Phaser^47^ using: UbcH8 structure (PDB 1WZW) as the E2 search model, and ISG15 (PDB 7S6P) as the substrate search model. Iterative cycles of model building using Coot^48^, and refinement using REFMAC5^49^ were performed. The final atomic model was validated using MolProbity^50^ and deposited in the Protein Data Bank (accession 9LW4). All structural refinement parameters fell within expected ranges for the resolution (Extended Data Table 1).

AlphaFold3^51^ predictions were generated for protein complexes as indicated in the figures, using default parameters with no constraints. The predicted model with the highest IDDT score was used in the final analysis, and PAE scores for various models are shown in Extended Data Figure 7. The model of UbcH8∼ISG15 and HERC5 complex in Figure 5 was generated by superposition of UbcH8/ISG15/HERC5 on the cryo-EM structure of UbcH5/Ub/ Pub2 (HECT) (PDB 9B55).

### Statistical Analysis and Data Quantification

All quantitative data represent triplicates (n=3 independent experiments). Data visualization and statistical analyses were performed using GraphPad Prism version 9.0. Statistical significance was determined using an unpaired t-test for comparisons between two groups, and one-way or two-way ANOVA for comparisons among multiple groups; significance was defined as \***p < 0.001.** Experimental variability is presented as mean ± standard error of the mean (SEM) in all graphical representations.

## Extended Data Figures

**Extended Data Figure 1 | Distinct conformations of E2∼Ub complexes with different E3 ligase families.**

**a** Crystal structure of UbcH5A∼Ub in complex with the RING domain of RNF4 (PDB 4AP4). The catalytic cysteine of UbcH5A is marked with a yellow sphere. Domain boundaries of RNF4 are indicated below.

**b** Crystal structure of UbcH7∼Ub bound to the RBR domain of HOIL1 (PDB 8EAZ). The catalytic cysteine of UbcH7 is highlighted (yellow sphere). Domain boundaries of HOIL1 are shown below.

**c** Crystal structure of UbcH5B∼Ub in complex with the HECT domain of NEDD4 (PDB 3JW0). The catalytic cysteine of UbcH5B is shown as a yellow sphere. Domain boundaries of NEDD4 are annotated.

**Extended Data Figure 2 | Activity assay using different family E3 ligases.**

**a** Ubiquitination assay using E3 ligases from different families. IR^800^ scan showing ubiquitination activity; an ATP-independent background band is marked with an asterisk (*). **b** ISGylation assay using WT/C867A NEDD4 E3 ligase. IR^800^ scan showing ISGylation activity (top panel); an ATP-independent background band is marked with an asterisk (*).

**Extended Data Figure 3 | UbcH8∼cISG15 complex preparation and structural analysis. a** Synthesis and purification of the UbcH8∼cISG15 complex. Left, schematic representation of the chemical strategy used to stabilize the UbcH8∼cISG15 complex. Right, size-exclusion chromatography (SEC) profile showing separation of the UbcH8∼cISG15 complex (fraction 1) from unreacted components (fractions 2 and 3). SDS-PAGE analysis confirms the identity of the eluted fractions.

**b** The electron density (2Fo-Fc) map (grey) for ISG15 (pink) and UbcH8 (cyan) around the catalytic pocket of UbcH8∼cISG15 crystal structure, contoured at 1.2 σ.

**c** Structural comparison of cryo-EM structures of UBE1-Ub(a)-Cdc34-Ub(t) (left) and UBE1L-ISG15(a)-UbcH8-ISG15(t) (right). The positions of the I44 (Ub) and T125 (ISG15) residues are highlighted in red and blue, respectively.

**d** E2 charging assay comparing different UbcH8 variants (upper panel). Lower panel, Coomassie-stained gel used as a loading control.

**e** E2 charging assay comparing WT or W123R/P130Q ISG15 (upper panel). Lower panel, Coomassie-stained gel used as a loading control.

**Extended Data Figure 4 | Functional analysis of predicted model of UbcH8∼ISG15/HERC5 complex.**

**a** Quantification of ISGylation activity of HERC5 using UbcH8 variants relative to WT UbcH8. Left, representative IR^800^ scan of the assay used for quantification. A non-specific band is marked with an asterisk (*); Coomassie-stained gel (bottom panel) serves as a loading control.

**b** E2 charging assay comparing activity of different UbcH8 variants using non-reducing (- DTT) SDS-PAGE (upper panel) and reducing (+DTT) SDS-PAGE (middle panel). Coomassie-stained gel (bottom) serves as a loading control.

**Extended Data Figure 5 | Functional analysis of the ISG15 donor-binding site on HERC5.**

**a** In vitro ubiquitination assay measuring HERC5 activity using the various E2 ubiquitin conjugating enzymes. IR^800^ scan shows ubiquitin conjugation; non-specific bands are marked with an asterisk (*). Coomassie-stained gel showing E2 (lower panel) is used as a loading control.

**b** Ubiquitination assay using NEDD4 variants to evaluate the functional relevance of specific residues (top). Bottom, Coomassie-stained gel provides a loading control.

**c** E2 discharge assays assessing the rate of E2-to-E3 ISG15 transfer mediated by wild-type or P988L/E1015M mutant HERC5. Representative IR^800^ scan showing discharge at indicated time points (top). Quantification from three biological replicates (bottom).

**Extended Data Figure 6 | Sequence alignment of UbcH8 with other E2 conjugating enzymes.**

Multiple sequence alignment of UbcH8 with representative E2 ubiquitin-conjugating enzymes. Secondary structure elements are shown above the alignment. Residues known to mediate E2–E3 interactions are indicated with arrows.

**Extended Data Figure 7 | PAE scores of models of HERC5∼ISG15 (a), HERC5/UbcH8 (b), and NEDD4/UbcH6 (c) predicted using Alphafold**

## Authors’ contributions

P.S. conducted all biochemical/biophysical experiments and performed data analysis. G.G.P. purified, crystallized, and analyzed the UbcH8∼cISG15 complex. D.R.L., M.S., and B.S.T. provided essential reagents. D.R.L. additionally contributed to X-ray data collection and processing. A.K. conceived the project, designed the study, supervised the research, determined structures, analyzed data, and acquired funding. A.K. wrote the manuscript with critical input from P.S. and G.G.P. All authors reviewed and approved the final manuscript.

## Acknowledgements

The authors are grateful to the anonymous reviewers for their insightful comments that strengthened this work. We also acknowledge Prof. Deepak Nair (Regional Centre for Biotechnology, Faridabad) and the Department of Biotechnology (Government of India) for facilitating access to ESRF beamtime. Special thanks to Dr. Shaun K. Olsen and Dr. Digant Nayak (University of Texas Health Science Center at San Antonio) for their valuable guidance on UBE1L expression in insect cells. We thank the Central Instrumentation Facility at IISER Bhopal for providing access to the ITC instrument. We also thank members of the Kumar lab for their constructive feedback on the manuscript and reagent support. P.S., G.G.P., and M.S. acknowledge Senior Research Fellowship support from the Council of Scientific and Industrial Research (CSIR). D.R.L. is supported by the Prime Minister’s Research Fellowship (PMRF). A.K. is a recipient of the DBT-Ramalingaswami Re-entry Fellowship (BT/RLF/Re-entry/42/2019) and acknowledges research funding from IISER Bhopal, Department of Biotechnology (BT/12/IYBA/2019/03).

