## Supplementary for "ISGylation Mechanism Uncovers Conformational Specificity for HECT-family E3 ligase": ExtendedDataFig_Merged.pdf

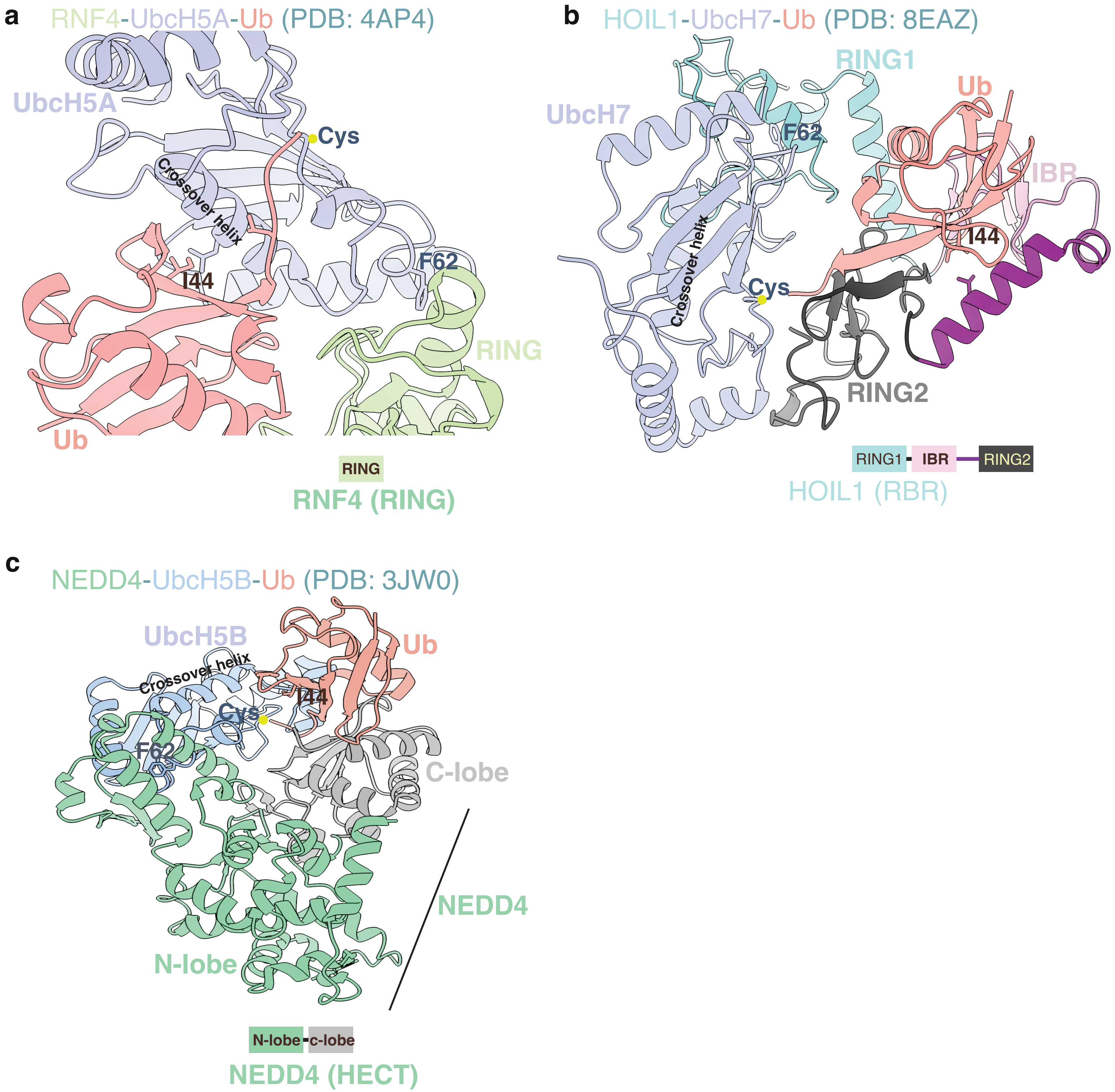
a
RNF4-UbcH5A-Ub (PDB: 4AP4)
UbcH5A
Cys
Crossover helix
I44
F62
RING
Ub
RING
RNF4 (RING)
b
HOIL1-UbcH7-Ub (PDB: 8EAZ)
RING1
Ub
UbcH7
F62
IBR
Crossover helix
I44
Cys
RING2
RING1
IBR
RING2
HOIL1 (RBR)
c
NEDD4-UbcH5B-Ub (PDB: 3JW0)
UbcH5B
Ub
Crossover helix
I44
Cys
F62
C-lobe
NEDD4
N-lobe
N-lobe
c-lobe
NEDD4 (HECT)

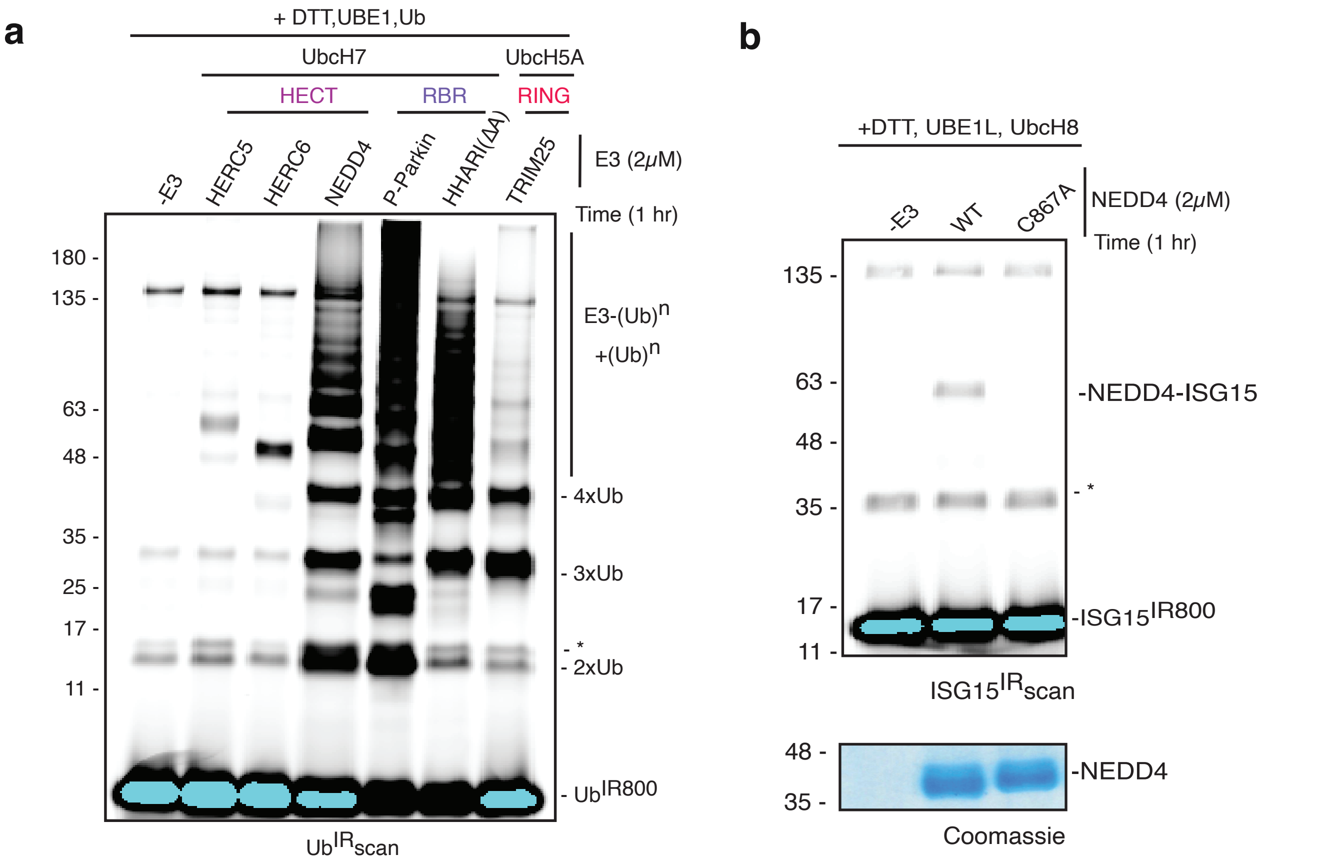


**Extended Data Figure 2**

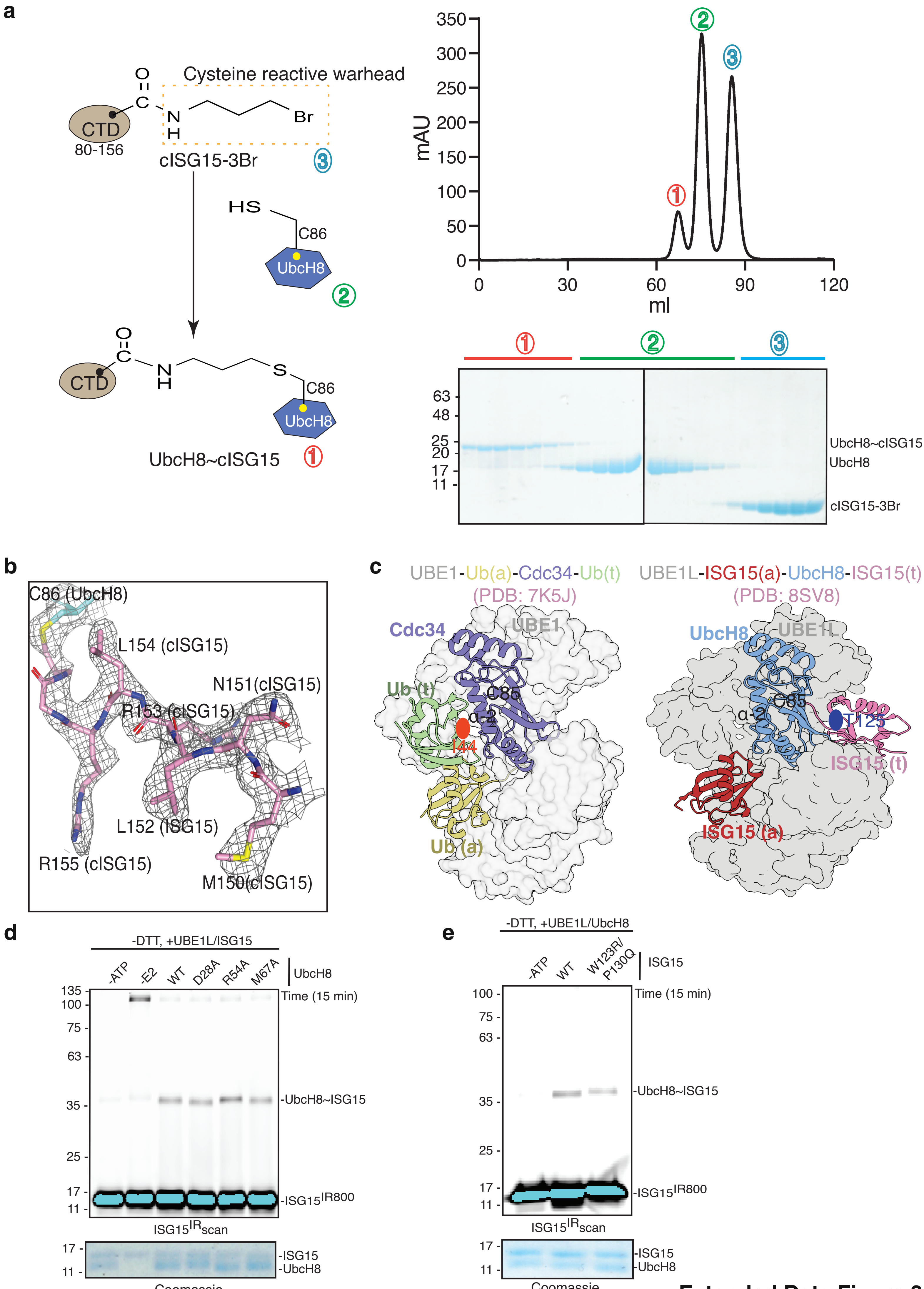


**Extended Data Figure 3**

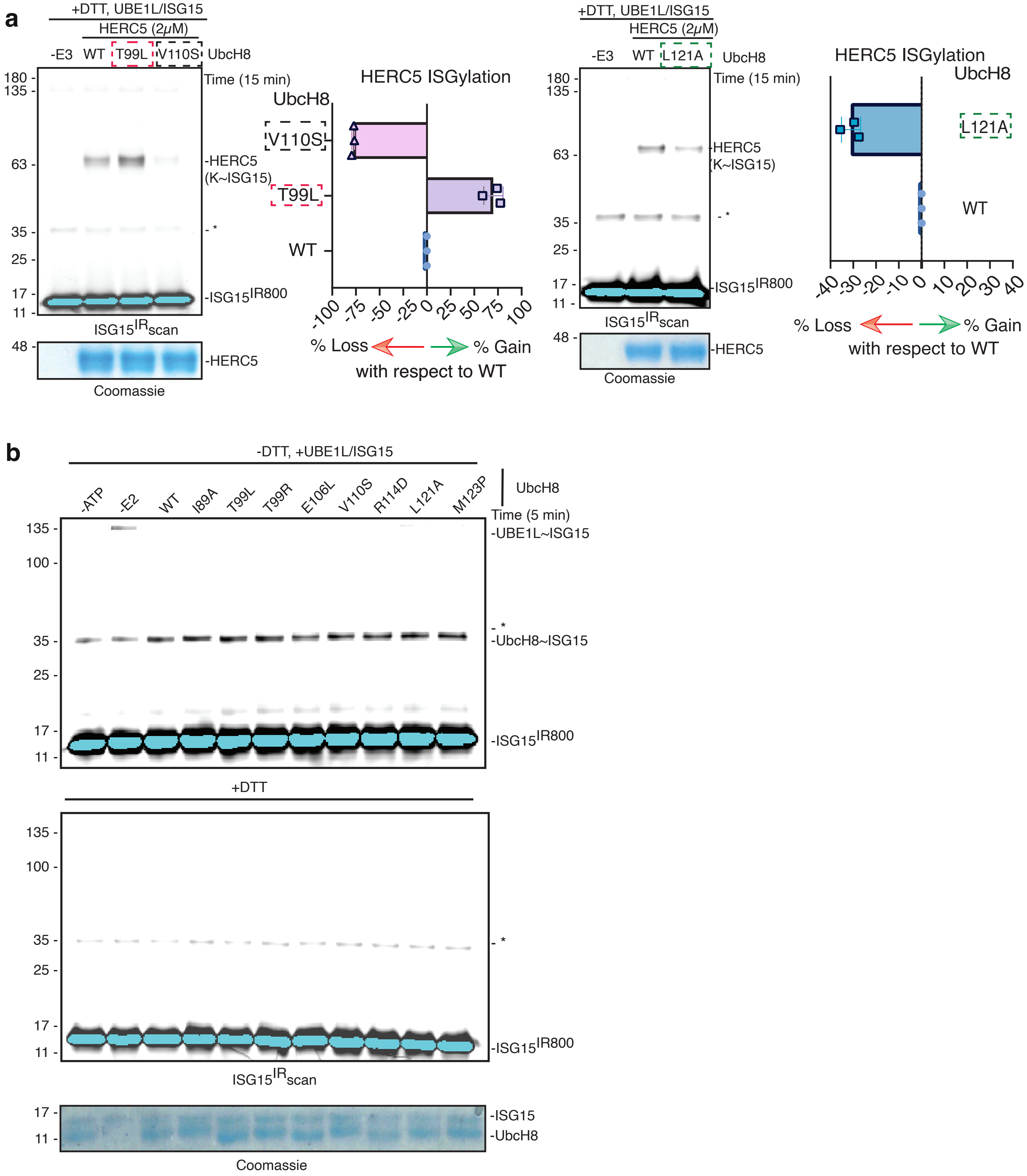


**Extended Data Figure 4**

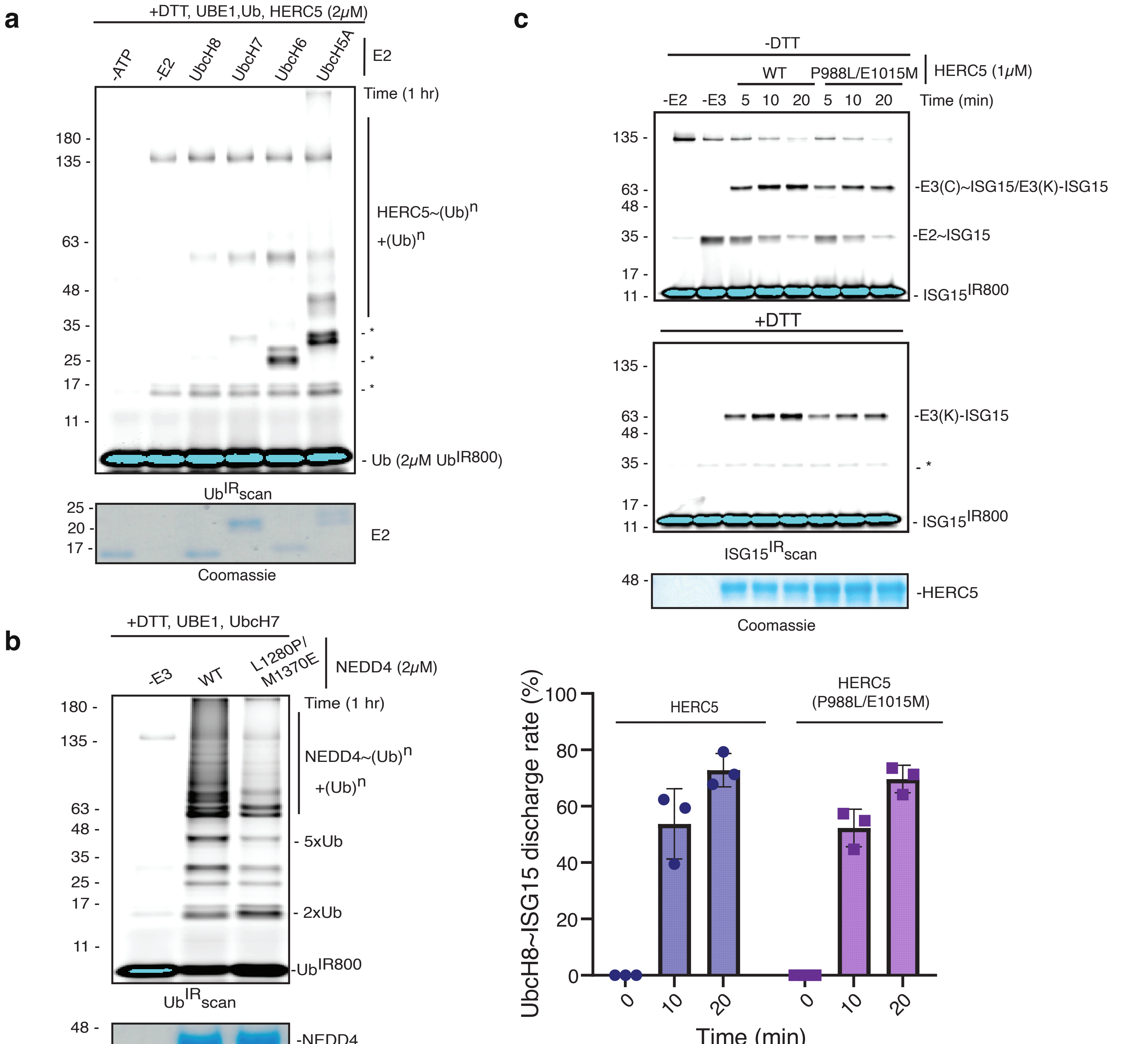


**Extended Data Figure 5**

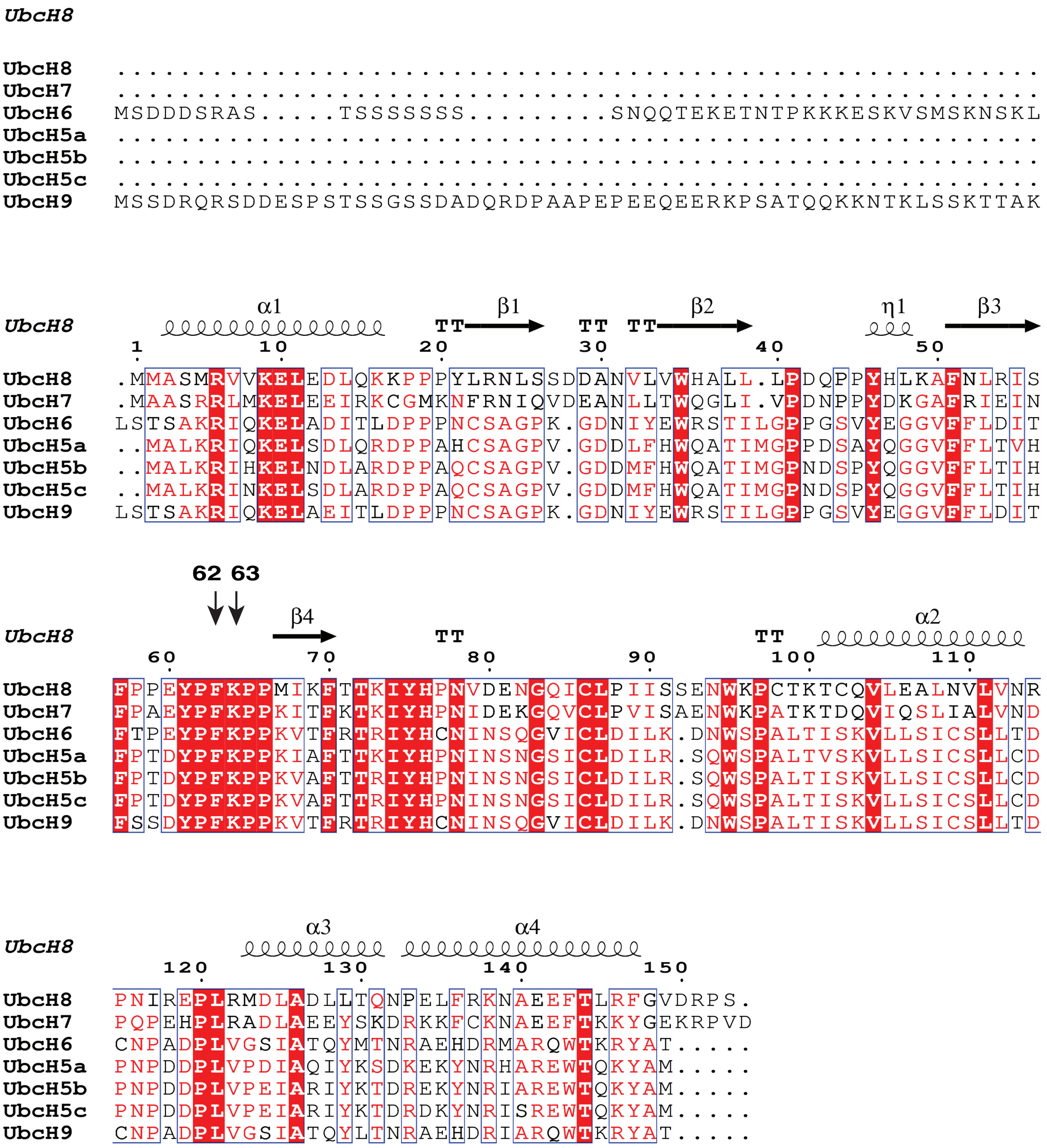


**Extended Data Figure 6**

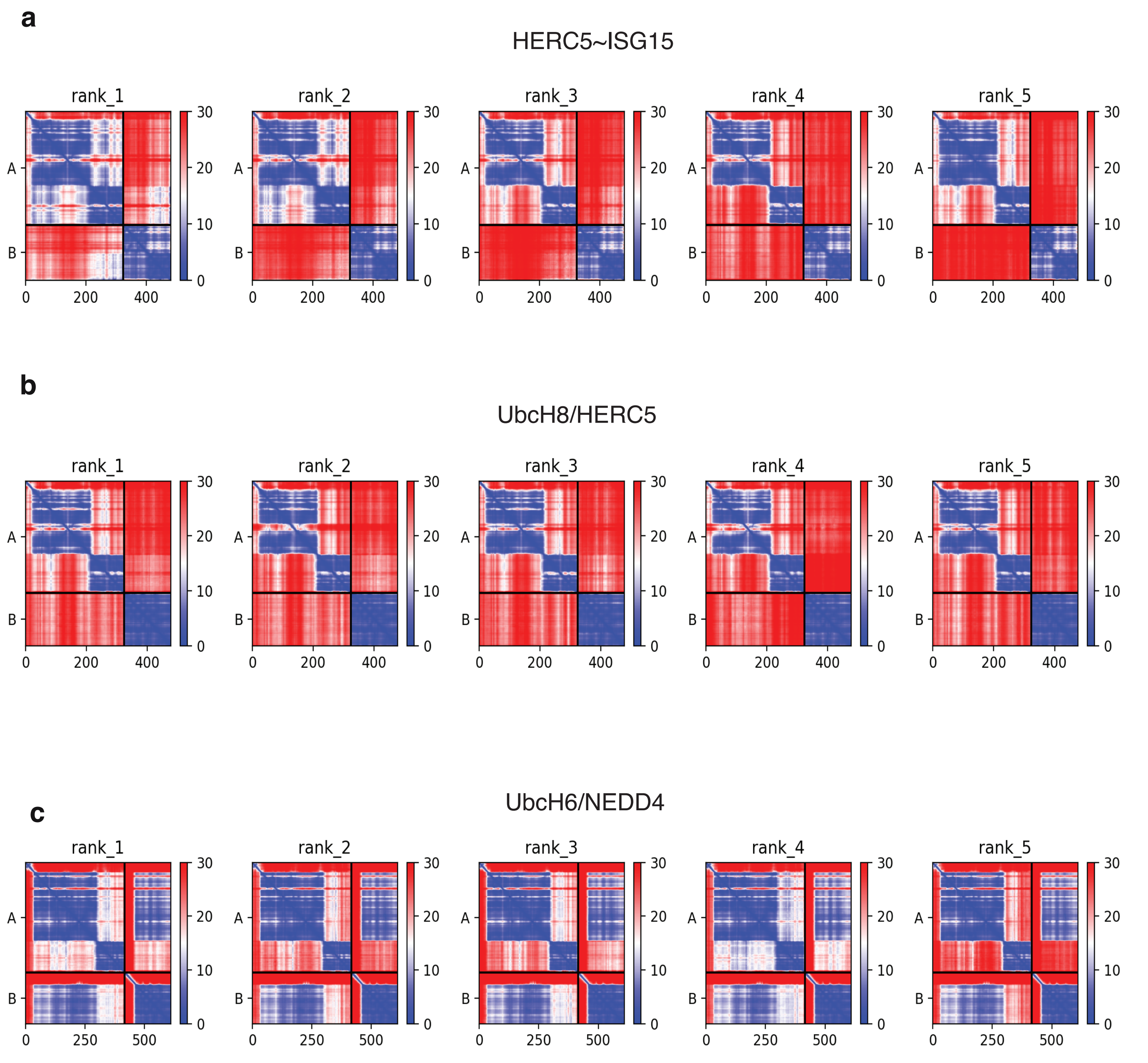


**Extended Data Figure 7**
